# Strong sustained type I IFN signaling acts cell intrinsically to impair IFNγ responses and cause tuberculosis susceptibility

**DOI:** 10.64898/2026.01.19.700487

**Authors:** Stefan A. Fattinger, Bianca Parisi, Roberto A. Chavez, Marian R. Fairgrieve, Ophelia V. Lee, Kristen C. Witt, Jesse J. Rodriguez, Elizabeth A. Turcotte, Ella C. Brydon, Harmandeep Dhaliwal, Angus Y. Lee, Dmitri I. Kotov, Russell E. Vance

## Abstract

*Mycobacterium tuberculosis* (*Mtb*) causes over one million deaths annually, but most infected individuals never exhibit symptoms. Type I interferons (IFNs) have emerged as a major factor driving susceptibility to *Mtb*, but how type I IFNs impair immunity to *Mtb* is a key unresolved question. Here we show that an early effect of type I IFN during *Mtb* infection is the cell-intrinsic impairment of IFNγ signaling. IFNγ signaling was selectively impaired in the subset of infected macrophages experiencing high and sustained levels of type I IFN signaling. Genetic elimination of RESIST, a recently described positive regulator of type I IFN production, specifically eliminated the high and sustained type I IFN response, fully restored IFNγ signaling, and rescued susceptibility to *Mtb* without affecting basal type I IFN responses. Our results demonstrate that strong and sustained type I IFN responses specifically and cell intrinsically impair responsiveness to IFNγ to cause susceptibility to *Mtb*.

## Introduction

Tuberculosis (TB) is caused by *Mycobacterium tuberculosis* (*Mtb*) and is the deadliest infectious disease of humans, with over one million yearly deaths world-wide^1^. The standard treatment for TB involves a 4-6 month course of antibiotics that is often poorly tolerated and ineffective against increasingly prevalent multi-drug resistant *Mtb. Mtb* infection elicits a robust innate and adaptive immune response that protects most but not all infected individuals from disease. A major unresolved question is how variations in the immune response to *Mtb* result in severe disease in some individuals. Interferon-γ (IFNγ) is implicated in protection against mycobacterial infections in humans^2–5^, and is essential in mice for resistance to *Mtb*^6–10^. However, most susceptible humans and mice exhibit robust IFNγ production in response to *Mtb* infection, indicating that IFNγ is often insufficient for protection. Indeed, BCG and a recent vaccine candidate that induced IFNγ-production by *Mtb*-specific T cells nevertheless fail to protect against infection in adults^11–15^. In contrast to IFNγ, type I IFNs—including IFNβ, IFNα and other isoforms, all of which signal through the interferon-α/β receptor (IFNAR)—are clearly associated with progression of *Mtb* disease in both mice and humans^16–30^. A consistent finding is that type I IFN signaling impairs IL-1-mediated immunity by various mechanisms, including by modulating lipid mediator production, or by the upregulation of IL-1 receptor antagonist (IL-1Ra) or IL-10^25,29,31^. However, type I IFN signaling can promote susceptibility even in the absence of IL-1 signaling^25^, and IL-1Ra-deficiency also rescues mouse models lacking type I IFN-driven susceptibility^32^. Thus, existing data suggest that additional mechanisms, beyond suppression of IL-1, may contribute to type I IFN–driven susceptibility^33,34^.

Recently, we identified SP140 as a key negative regulator of type I IFN responses^35,36^. Loss of SP140 results in increased production of type I IFNs, uncontrolled *Mtb* replication, hypoxic granulomas, and an exacerbated inflammatory phenotype similar to that seen in humans with clinical TB^27,35,37^. The susceptibility of *Sp140*^−/−^ mice to *Mtb* is rescued by crosses to *Ifnar*^−/−^ mice, establishing *Sp140*^−/−^ mice as a useful model of type I IFN-driven tuberculosis disease. *Ifnar*-deficiency does not greatly impact the susceptibility of wild-type B6 mice to *Mtb*, implying that the basal levels of type I IFNs seen in B6 mice are not pathogenic during *Mtb* infection^29,31,38–40^. A major outstanding question is how elevated, but not basal, levels of type I IFN specifically impair immunity to *Mtb*.

SP140 is a transcriptional repressor that represses two tandemly duplicated genes, *Resist1* and *Resist2*, both of which encode the identical protein RESIST (REgulated Stimulator of Interferon via Stabilization of Transcript)^36^. *Sp140*^−/−^ macrophages express higher levels of RESIST, which stabilizes *Ifnb1* mRNAs by specifically impairing the activity of the CCR4-NOT polyA-tail deadenylase to promote *Ifnb1* mRNA turnover^36^. *Resist1/2*-deficiency eliminates the elevated levels of *Ifnb1* mRNA and IFNβ protein seen in *Sp140*^−/−^ mice, and returns IFNβ expression to the basal levels seen in B6 mice^36^, but whether this also restores resistance to *Mtb* remains unknown.

In the lungs of humans and mice, intracellular *Mtb* replication occurs mainly within a heterogenous population of CD64^+^ macrophages including interstitial macrophages (IMs), a type of myeloid cell in which T cells are only inconsistently able to mediate *Mtb* control^41,42^. Some replication may also occur in neutrophils^43,44^. Cell-type specific deletion of *Ifnar* in CD64^+^ myeloid cells rescues the susceptibility of *Sp140*^−/−^ mice^27^, implying that type I IFNs act on myeloid cells, but it remains unclear whether type I IFNs act cell intrinsically on infected macrophages to impair control of *Mtb* replication, or instead act cell extrinsically, e.g., by inducing macrophage production of cytokines such as IL-10 to suppress responses by other cells. We hypothesized that inconsistent IFNγ-mediated control of *Mtb* might arise from differential exposure of infected macrophages to type I IFNs. In support of this hypothesis, it has been shown that type I IFN signaling can impair the induction of certain IFNγ-induced interferon stimulated genes (ISGs)^45–48^. Moreover, elevated type I IFN ISG induction correlated with decreased expression of IFNγ-induced ISGs and detrimental outcomes in human *Mycobacterium leprae* infections^49^. Similar negative correlations between type I IFN responses and IFNγ responses have also been observed in *Listeria*-infected mice^50^. During *Mtb* infection, *in vitro* experiments have shown that type I IFN can impair the expression of IL-12—a key inducer of IFNγ production—in a cell-extrinsic manner by inducing IL-10^51^. *In vivo*, we recently found that the enhanced type I IFN response seen in *Sp140*^−/−^ mice correlates with an attenuated IFNγ response in *Mtb*-infected interstitial macrophages^27^. However, it was not shown that type I IFNs caused the decreased responsiveness to IFNγ. In addition, these analyses were performed at day 25 post infection, when bacterial loads had already diverged between susceptible and resistant mice, making it difficult to disentangle cause and effect. Moreover, there was no evidence that the cells exhibiting decreased IFNγ responsiveness were a susceptible niche for *Mtb* replication. Thus, it remains unclear if there is a causal relationship in which type I IFN signaling cell intrinsically impairs the IFNγ response of infected macrophages *in vivo*, and whether such impairment causes susceptibility to *Mtb*.

Here, we take advantage of *Sp140*^*–/–*^ mice to systematically dissect the emergence and cause of type I IFN-driven susceptibility to *Mtb*. We show that strong and sustained type I IFN signaling selectively impairs responses to IFNγ by a subset of IMs *in vivo* during *Mtb* infection. This subset of IMs also exhibits the highest burdens of *Mtb*. We further uncover a temporal progression in which type I IFN signaling precedes and subsequently diminishes IFNγ responsiveness, thereby rendering IMs a permissive intracellular niche for *Mtb*. Finally, by genetic elimination of RESIST in *Sp140*^−/−^ mice, we demonstrate that sustained and strong (but not basal) type I IFN signaling specifically suppresses IFNγ responsiveness and drives susceptibility to *Mtb*.

## Results

### IFNAR-STAT2-IRF9 signaling broadly impairs IFNγ signaling

To test whether type I IFN globally impairs the IFNγ-induced transcriptional responses, or just affects certain IFNγ-induced genes as previously shown^45–48^, we performed RNAseq on mouse bone marrow-derived macrophages (BMMs) exposed to IFNβ (type I IFN) alone, IFNγ alone, or both cytokines simultaneously (conditions referred to as β, γ and β/γ, respectively; Figure 1A). Most ISGs are similarly induced by IFNβ and IFNγ, but we previously defined a subset of 21 “IFNγ signature genes” that are preferentially induced by IFNγ and that are upregulated during *Mtb* infection^27^. Pre-exposing BMMs to IFNβ blunted IFNγ-induced expression of almost all IFNγ signature genes (Figure 1B). A notable exception was *Gbp8*, which was induced by IFNγ similarly in IFNβ-pretreated and control cells. To test if type I IFN signaling impairs the IFNγ response beyond these previously defined IFNγ signature genes, we defined a broader subset of 121 ISGs that are induced at least two-fold more by IFNγ as compared to IFNβ (Figure 1C). Strikingly, 91% (110/121) of these ISGs were suppressed by pre-exposure to IFNβ (Figure 1D). *Gbp2b, Gbp8* and *Gbp10* were among the 11 ISGs (Table S1A) that were induced by IFNγ despite IFNβ pretreatment. The chemokine CXCL9 is a canonical IFNγ-induced ISG that is highly expressed during human and non-human primate *Mtb* infections, and has been suggested to promote an effective T cell mediated immune response^52,53^. In our RNAseq dataset, *Cxcl9* was one of the most IFNγ-upregulated gene and was induced >8-fold more by IFNγ than IFNβ (Figure 1C), a result that we validated by RT-qPCR (Figure S1B). Strikingly, despite some induction of *Cxcl9* transcripts by IFNβ (Figures 1C, S1B), intracellular staining and flow cytometry showed that IFNβ pre-exposure fully abolished the induction of CXCL9 protein by IFNγ (Figures 1E, S1C). Exposure to IFNβ, IFNγ, or both did not impact cell viability (Figure S1D), and as expected, IFNβ suppression of CXCL9 induction required IFNAR (Figures 1E, S1C). While a 10-fold increase in IFNγ concentration could not overcome the IFNβ-mediated suppression of CXCL9 induction by IFNγ, co-stimulation with TNF or TLR-agonists partially reversed the inhibition in a IFNβ concentration-dependent manner (Figure S1F). The impairment of IFNγ signaling persisted after the removal of IFNβ, at a timepoint when type I IFN specific *Rsad2* transcripts (encoding Viperin) had already returned to baseline (Figures 1F, S1G, compare β/γ to β→γ, see Figure 1A for exposure details). Indeed, a time course analysis revealed that impairment of IFNγ signaling can last up to 32h upon IFNβ removal (Figure S1H).

**Figure 1.**
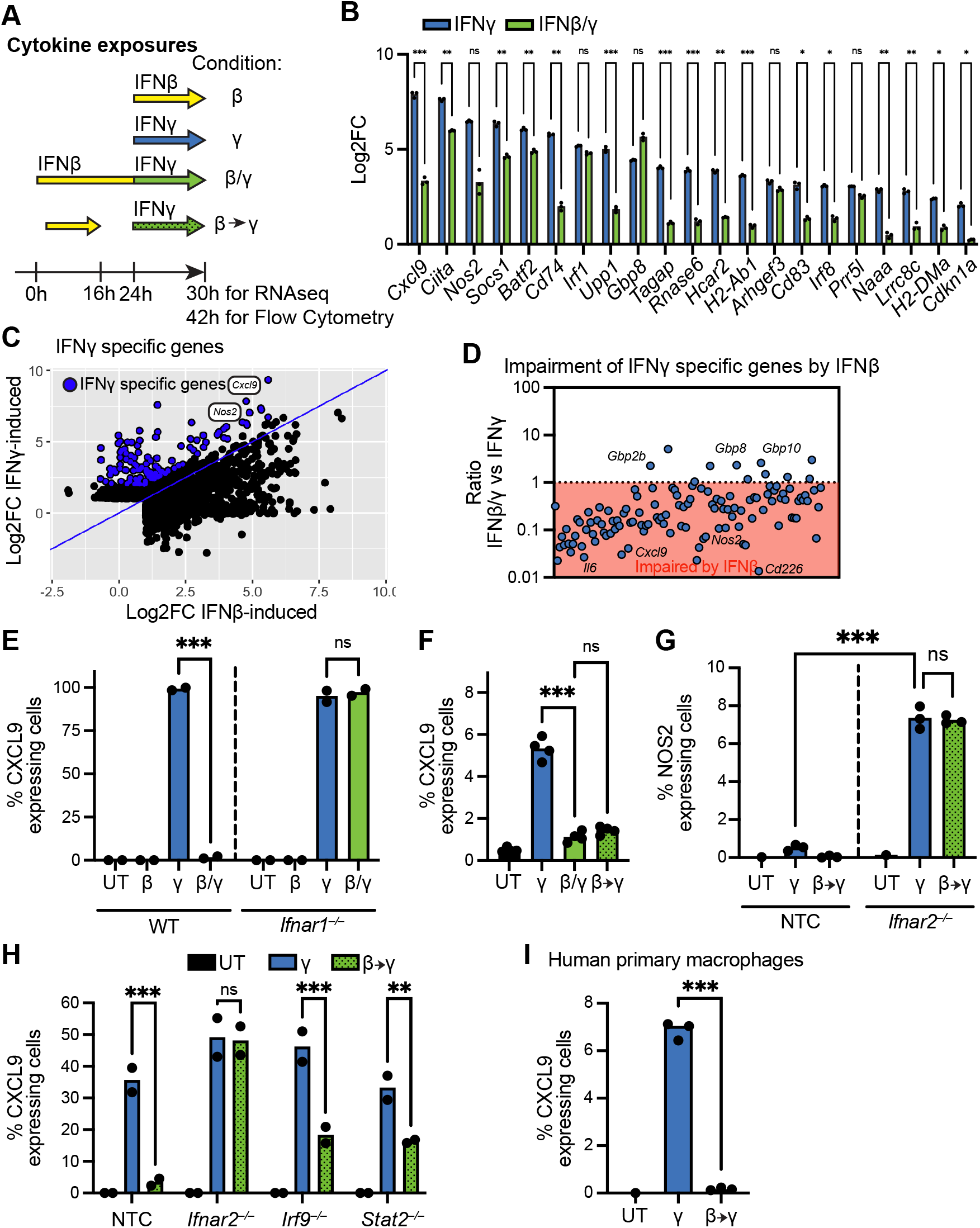
IFNAR-STAT2-IRF9 signaling broadly impairs IFNγ signaling. (**A**) Cytokine exposures for RNAseq and flow cytometry analysis. (**B**) Log_2_ fold change of IFNγ signature genes in bone marrow derived macrophages (BMMs) upon exposure to IFNγ ± IFNβ pre-exposure (*n* = 3). (**C**) Induction of IFN-specific genes by BMMs upon IFNγ and IFNβ. Genes two-fold more induced upon IFNγ compared to IFNβ are defined as IFNγ specific genes (blue). (**D**) Ratio of IFNγ-specific gene induction in BMMs upon IFNβ/γ vs IFNγ treatment. Average of *n* = 3 is plotted in (C-D). (**E**) CXCL9 protein expression by BMMs from B6 wild-type (WT) or *Ifnar1*^*–/–*^ mice upon exposure to IFNβ, IFNγ or IFNβ/γ. UT, Untreated (**F**) CXCL9 expression by BMMs, comparing IFNβ pre-exposure without (β/γ) or with (β→γ) its removal prior to IFNγ stimulation. (**G**) NOS2 expression by CRISPR-edited BMMs. NTC, Non-Target Control. (**H**) CXCL9-expression by CRISPR-edited BMMs. NTC, Non-Target Control. (**I**) CXCL9 expression by human primary macrophages. (E-I) Representative data from ≥ 3 biological replica with n ≥ 2 per condition. Statistical significance was calculated in (B) with RM two-way ANOVA with the Geisser-Greenhouse correction and Sidak’s multiple comparison test; in (E,G,H) with two-way ANOVA with (E,G) Sidak’s multiple comparison test or (H) Turkey’s multiple comparison test; in (F,I) one-way ANOVA with Sidak’s multiple comparison test. *p < 0.05, **p < 0.01, ***p < 0.001, ns = not significant.

Nitric Oxide Synthase 2 (NOS2) is another IFNγ-induced ISG implicated in *Mtb* protection in mice^8,54–56^. In our RNAseq dataset, *Nos2* transcripts were also highly and specifically expressed upon IFNγ (Figure 1C). In sharp contrast to CXCL9, NOS2 was barely detectable at the protein level when cells were treated with IFNγ (Figure 1G) consistent with previous reports^57^. Strikingly, however, *Ifnar2*^−/−^ BMMs generated by CRISPR induced abundant NOS2 protein in response to IFNγ (Figure 1G). This result implies that tonic type I IFN-signaling is sufficient to impair IFNγ-induced NOS2 expression and explains the previously observed failure of IFNγ to induce NOS2 in wild-type cells^57^. However, unlike CXCL9, NOS2 protein expression was not induced exclusively by IFNγ, and TLR agonists also fully reversed its suppression by IFNβ (Figure S1I). Given these observations, and the fact that the role of NOS2 in protection from TB in humans is still debated^58^, we chose CXCL9 as the most reliable and specific marker for the IFNγ response in subsequent analyses.

Signaling downstream of IFNAR involves the transcription factors STAT1, STAT2 and IRF9 that are believed to form a complex called Interferon-stimulated gene factor 3 (ISGF3) that binds interferon-sensitive response elements (ISREs) to induce ISG expression^59^. To determine whether these signal transducers are involved in the impairment of IFNγ responsiveness, we generated STAT2 and IRF9 deficient BMMs, and then exposed the knockout cells to IFNγ with or without IFNβ pre-exposure. Both STAT2 and IRF9-deficiency partially prevented type I IFN-driven inhibition (Figure 1H).

Next, to confirm the relevance of our findings to human cells, we exposed human immortalized THP-1 cells and primary monocyte derived macrophages to IFNs. In both human-derived cell types, IFNβ robustly inhibited IFNγ-induced CXCL9 induction, demonstrating that type I IFN exposure also impairs the IFNγ response in human macrophages (Figures 1I, S1J). Together, these data establish CXCL9 protein as a marker for IFNγ signaling in macrophages, and along with prior observations^45–48^, demonstrate that in mouse and human macrophages, type I IFN signaling downstream of IFNAR can engage STAT2 and IRF9 to broadly impair responsiveness to IFNγ in a robust and sustained manner.

### Type I IFNs impair the IFNγ response to *Mtb in vivo*

*Sp140*^−/−^ mice express elevated levels of type I IFN and are a robust model of type I IFN driven susceptibility to *Mtb*^35^. We infected *Sp140*^−/−^, *Sp140*^−/−^*Ifnar1*^*–/–*^ and *Sp140*^−/−^*Ifnar1*^*fl/f*^ CD64^Cre^ mice for 25 days with a low aerosolized dose (20-100 colony forming units (CFUs)) of *Mtb* Erdman strain expressing a fluorescent reporter, the intensity of which was previously validated to correlate with *Mtb* CFU *in vivo*^27^. We confirmed that type I IFNs mainly act on CD64^+^ myeloid cells to cause susceptibility^27^ (Figure 2A). To determine if CD64^+^ myeloid cells are also the main cell type on which IFNγ acts to control *Mtb* replication, we generated *Ifngr1*^*fl/fl*^CD64^Cre^ mice which specifically lack IFNGR on myeloid cells. Strikingly, *Ifngr1*^*fl/fl*^CD64^Cre^ mice were as susceptible as global *Ifngr1*-deficient mice (Figure 2A). This suggests that CD64^+^ myeloid cells are the target cells for both type I IFN driven susceptibility and IFNγ-dependent protection. Thus, we focused our analysis on CD64^+^ IMs that are also a main intracellular niche for *Mtb* replication^42^ (see Figure S2A for gating strategy).

Based on our *in vitro* results (Figure 1), we sought to test whether CXCL9 is a reliable marker for IFNγ responsiveness on infected IMs *in vivo*. At day 25 post infection (pi), while intracellular CXCL9 expression was negligible in neutrophils (Figures S2B-C), IMs from wild-type B6 mice expressed significantly higher levels of CXCL9 compared to *Ifngr1*^*–/–*^ mice (Figure 2B). Differences in autofluorescence of IMs across genotypes could be excluded as a confounding factor (Figures S2D-E). Thus, we selected CXCL9 as a reliable marker of IFNγ responsiveness *in vivo* during *Mtb* infection.

**Figure 2.**
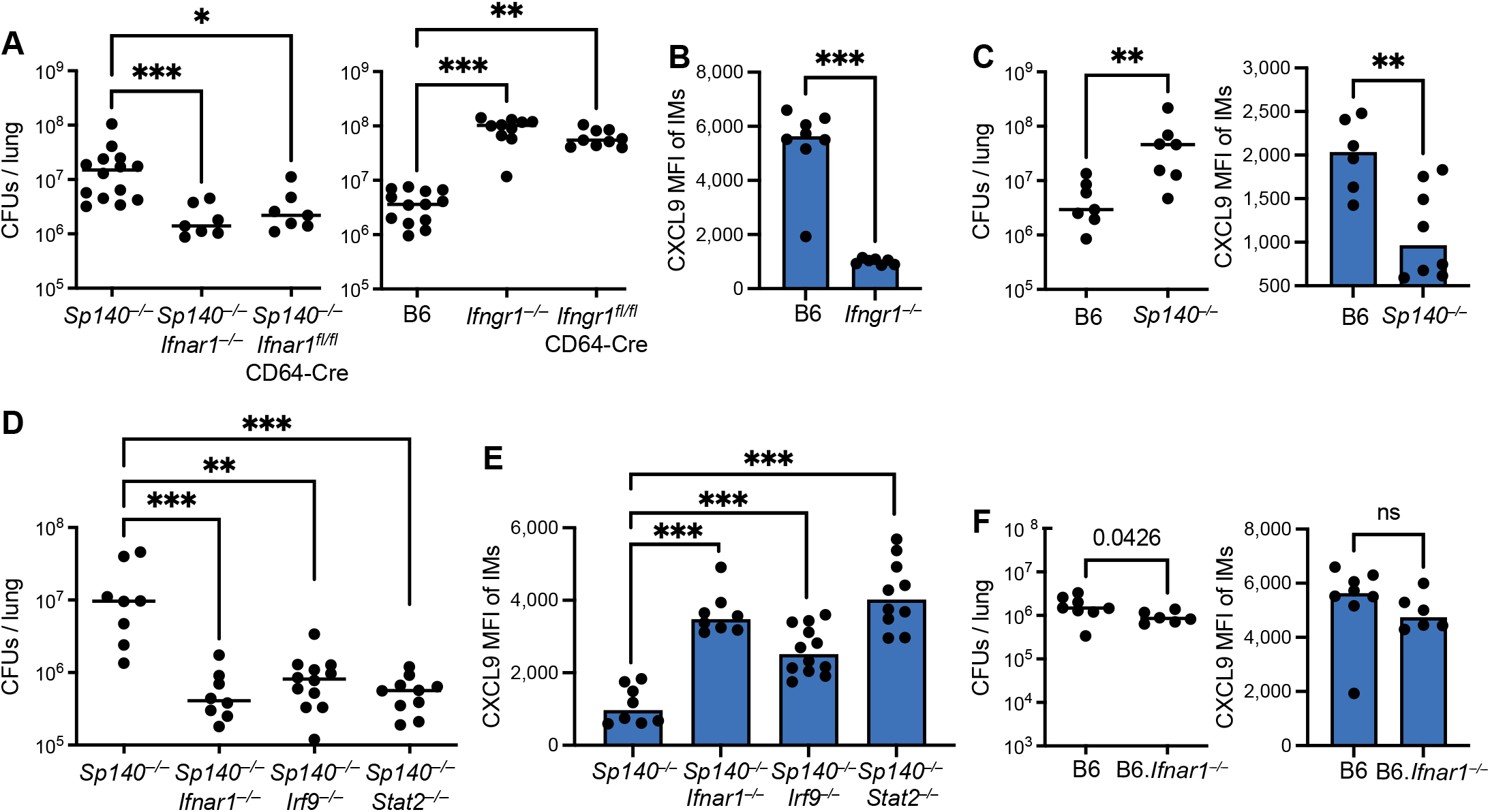
Type I IFNs impair the IFNγ response to *Mtb in vivo*. (**A**) Colony Forming Units (CFUs) per lung from *Mtb*-infected B6 or type I IFN susceptible *Sp140*^*–/–*^ mice with global or CD64^+^ macrophage-specific deletion of *Ifnar1* or *Ifngr1* at day 25 post infection (pi) (**B**) Mean Fluorescence Intensity (MFI) of CXCL9 staining of interstitial macrophages (IMs) from *Mtb*-infected B6 and *Ifngr1*^*–/–*^ mice at day 25 pi. (**C**) CFUs per lung and MFI of CXCL9 staining in IMs from *Mtb*-infected B6 and *Sp140*^*–/–*^ mice at day 25 pi. (**D**) CFUs per lung and (**E**) MFI of CXCL9 staining of IMs from indicated mouse strains at day 28-29 pi. (**F**) CFUs per lung and MFI of CXCL9 staining in IMs from *Mtb*-infected B6 and B6.*Ifnar1*^*–/–*^ mice at day 25-26 pi. In each figure, data was combined from ≥ 2 biological replica with n ≥ 6 per group. Statistical significance was calculated for CFU data in (A, D) with Kruskal-Wallis test with Dunn’s multiple comparison test in (C, F) with Mann-Whitney test and, for MFI data in (B,C,F) Welch’s t-test and in (E) Brown-Forsythe/Welch ANOVA test with Dunnett’s T3 multiple comparisons test. *p < 0.05, **p < 0.01, ***p < 0.001, ns = not significant.

Next, to test whether enhanced type I IFN signaling represses responsiveness to IFNγ *in vivo*, we infected B6 and *Sp140*-deficient mice. IMs from *Sp140*^−/−^ mice expressed lower levels of CXCL9, indicative of reduced IFNγ responsiveness, as compared to B6 mice (Figure 2C). To test whether the decreased expression of CXCL9 observed in *Sp140*^−/−^ mice was due to IFNAR signaling, we examined *Sp140*^−/−^*Ifnar1*^−/−^ mice. As previously reported^35^, IFNAR-deficiency rescued the susceptibility to *Mtb* seen in *Sp140*^−/−^ mice (Figure 2D). We also observed that IFNγ signaling in IMs (as read out by CXCL9 expression) is rescued in *Sp140*^−/−^*Ifnar1*^−/−^ mice as compared to *Sp140*^−/−^ mice (Figure 2E). Notably, *Sp140*^−/−^ mice harboring single *Stat2* or Irf9 null mutations (generated by CRISPR) were also rescued for type I IFN-driven susceptibility to *Mtb* and IFNγ responsiveness, as assessed by CXCL9 expression (Figures 2D, E). To validate these results, we also examined NOS2 expression. Compared to CXCL9, NOS2 was mainly expressed in infected (as opposed to uninfected bystander) IMs (Figures S2F-G). Focusing on infected IMs, we observed suppression of NOS2 expression in cells with intact type I IFN signaling (Figures S2H-I).

During *Mtb* infection, B6 mice express lower levels of type I IFN as compared to *Sp140*^*–/–*^ mice, and do not show a consistent type I IFN-driven susceptibility to *Mtb*^29,31,38–40^. Thus, if the susceptibility of *Sp140*^−/−^ mice is due to impairment of the IFNγ response by type I IFN signaling, we should not observe such impairment in wild-type B6 mice. Indeed, *Ifnar1*-deficiency had no effect on CXCL9 nor NOS2 expression on a B6 background (Figures 2F, S2J). If anything, we detected lower CXCL9 levels in B6.*Ifnar1*^*–/–*^ mice as compared to B6 IFNAR-sufficient mice.

Overall, our findings indicate that elevated and/or sustained levels of type I IFNs in *Sp140*^−/−^ mice impair the IFNγ response to *Mtb in vivo*, and that disruption of type I IFN signaling through STAT2 or IRF9 deficiency is sufficient to overcome this effect.

### Type I IFNs impair IFNγ-responsiveness and *Mtb* control cell intrinsically

Type I IFN signaling has been suggested to impair the IFNγ response cell intrinsically by downregulating IFNγ receptor levels, or cell extrinsically by upregulating the immunosuppressive cytokine IL-10^48–51^. To distinguish these possibilities, we mixed WT CD45.1 and *Ifnar1*^*–/–*^ CD45.2 BMMs at a ratio of 9:1 and exposed them to IFNβ and IFNγ (IFNβ/γ) *in vitro*. IFNβ impaired the response to IFNγ in WT cells, but *Ifnar1*^*–/–*^ cells in the same culture robustly induced CXCL9, indicating that IFNβ-responsive cells do not suppress neighboring non-responsive cells (Figure 3A). Next, we sought to identify an ISG that marks type I IFN–responsive cells by flow cytometry. We found that *Rsad2* (encoding Viperin) was abundantly expressed upon IFNβ exposure at the mRNA (Figure S1G) and protein levels (Figure 3B). Expression of Viperin protein was specific to type I IFN signaling (and not observed upon IFNγ exposure) and was dependent on IFNAR (Figure 3B). In mixed WT and *Ifnar1*^*–/–*^ BMM cultures exposed to IFNβ/γ, CXCL9 and Viperin expression were mutually exclusive, further supporting a cell-intrinsic mechanism for the type I IFN dependent impairment of the IFNγ responses (Figure 3C). Together, these data indicate that, *in vitro*, type I IFN acts cell intrinsically and does not induce a soluble mediator sufficient to suppress IFNγ signaling.

**Figure 3.**
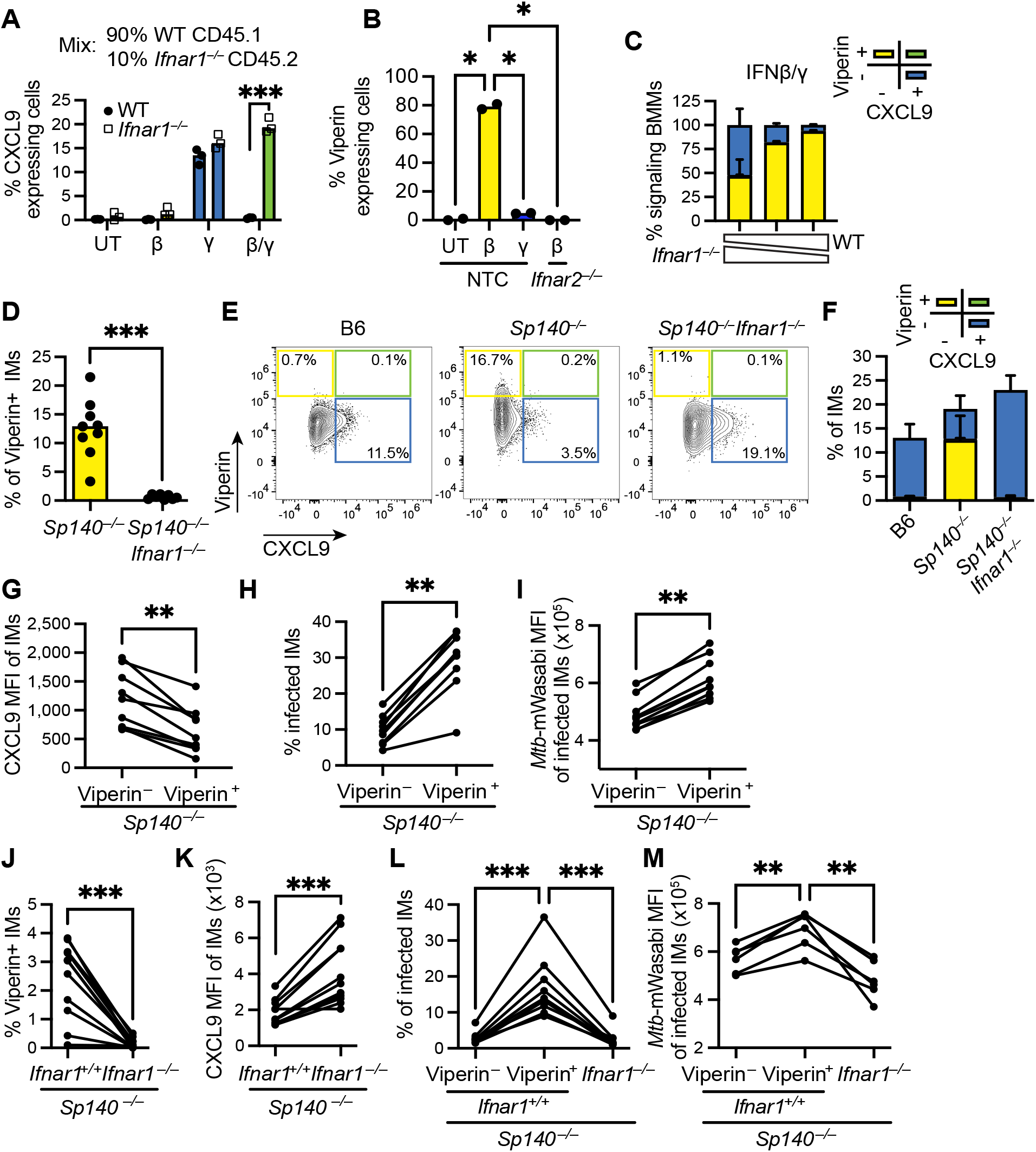
Type I IFNs impair IFNγ-responsiveness and *Mtb* control cell intrinsically. (**A**) Quantification of CXCL9 expression in WT CD45.1 and *Ifnar1*^*–/–*^ CD45.2 BMMs from a mixed culture with an overabundance of WT CD45.1 cells (9:1) upon exposure to IFN**γ**, IFNβ and both (IFNβ/**γ**). (**B**) Viperin-expression by CRISPR-edited BMMs upon IFN**γ** or IFNβ. NTC, Non-Targeting Control; UT, Untreated. (**C**) Quantification of CXCL9 and/or Viperin expressing cells from mixed WT : *Ifnar1*^*–/–*^ BMM cultures at various ratios upon exposure to both IFNβ and IFN**γ** (IFNβ/**γ**). (**D**) Percent Viperin expressing IMs from *Mtb*-infected *Sp140*^*–/–*^ and *Sp140*^*–/–*^*Ifnar1*^*–/–*^ mice at day 28-29 pi. (**E**) Representative flow cytometry plots and (**F**) analysis of IMs for CXCL9 and Viperin expression from *Mtb*-infected B6, *Sp140*^*–/–*^ and *Sp140*^*–/–*^*Ifnar1*^*–/–*^ mice at day 28-29 pi. (**G**) CXCL9 MFI, (**H**) percent *Mtb* infection, and (**I**) *Mtb*-mWasabi MFI in Viperin-positive and -negative IMs from *Mtb*-infected *Sp140*^*–/–*^ mice at day 28-29 pi. (**J-M**) *Mtb*-infected mixed *Sp140*^*–/–*^ bone marrow chimeras containing IFNAR-proficient (CD45.1) and -deficient cells (CD45.2) at day 25-26 pi analyzed for (**J**) percent Viperin positive IMs, (**K**) CXCL9 MFI, (**L**) percent *Mtb* infection, and (**M**) MFI of *Mtb*-mWasabi among Viperin/IFNAR-expressing IM subsets. (A-C) Representative data from ≥ 3 biological replica with n ≥ 2 per condition. (C,F) Data are represented as mean ±SD. (D, F-M) In each figure, data was combined from ≥ 2 biological replica with n ≥ 7 per group. Statistical significance was calculated (A-B) with Two-way ANOVA and Sidak’s multiple comparison test, in (D) Welch’s t-test, in (G-K) Paired t-test, in (L-M) RM one-way ANOVA with Sidak’s multiple comparison test. *p < 0.05, **p < 0.01, ***p < 0.001, ns = not significant.

We then examined Viperin and CXCL9 expression by IMs in *Mtb* infected mice. As expected, Viperin staining was IFNAR-dependent in *Mtb*-infected *Sp140*^*–/–*^ mice (Figure 3D) and not impacted by autofluorescence (Figures S3A-B). In line with the *in vitro* results, expression of CXCL9 and Viperin in IMs was mutually exclusive (Figures 3E-F), as expected if type I IFN responsive cells fail to respond to IFNγ. Moreover, in *Sp140*^−/−^ mice, Viperin-positive IMs not only expressed reduced levels of CXCL9 but were also more frequently infected and had elevated levels of *Mtb* (Figures 3G-I). These data suggested a direct connection between elevated type I IFN-signaling, reduced IFNγ-responsiveness, and elevated *Mtb* burdens within the same cell. However, a caveat of these experiments is that the results could be confounded by the different *Mtb* burdens in *Sp140*^−/−^ versus *Sp140*^−/−^*Ifnar1*^−/−^ mice.

To control for the different bacterial burdens of *Sp140*^*–/–*^ and *Sp140*^*–/–*^*Ifnar1*^*–/–*^ mice, we established mixed bone marrow chimeras in which CD45.1 IFNAR-proficient and CD45.2 IFNAR-deficient *Sp140*^−/−^ bone marrow cells were used to reconstitute *Sp140*^−/−^ recipients. We also generated control chimeras reconstituted with a mix of CD45.1 and CD45.2 bone marrow cells, both of which were IFNAR-proficient (Figures S3C-F). Mice were infected with *Mtb* >8 weeks after hematopoietic reconstitution. In these chimeras, IFNAR-proficient and IFNAR-deficient cells respond to *Mtb* and IFNγ in the same inflammatory environment. In line with the results above, Viperin expression was dependent on IFNAR signaling and CXCL9 expression was restored in *Ifnar1*^*−/−*^ IMs (Figures 3J-K). Notably, Viperin staining revealed that only a minor fraction (~1-4%) of *Ifnar1*^*+/+*^ IMs actively respond to type I IFN (Figure 3J, control in Figure S3C). Due to the transient induction of Viperin (Figure S1G), Viperin expression may fail to identify all the cells that have sensed type I IFNs (see below). Nevertheless, we consistently found decreased CXCL9 levels (Figure S3G) and elevated *Mtb* burdens in the few Viperin expressing *Ifnar1*^*+/+*^ IMs compared to Viperin-negative *Ifnar1*^*+/+*^ cells or to *Ifnar1*^−/−^ IMs (Figures 3L-M). Although some IFNAR^+^ but Viperin-negative cells may never have been exposed to IFNβ, it was still possible to detect increased CXCL9 expression and decreased *Mtb* infection by percentage and MFI in the total *Ifnar1*^*+/+*^ cell population (Figures S3D-F).

In summary, these observations suggest that in IMs, type I IFN predominantly impairs IFNγ response in a cell-intrinsic manner to promote susceptibility to *Mtb*.

### Sustained and strong type I IFN–signaling impairs IFNγ responsiveness

Although Viperin expression marked some type I IFN-responsive cells, Viperin expression is transient and is lost before the repressive effects of type I IFN on IFNγ signaling have resolved (Figures 1F, S1G). Thus, Viperin expression likely underreports previous exposure to type I IFN. As a more sensitive reporter for type I IFN signaling, we took advantage of a previously described type I IFN signaling reporter mouse^60^ in which green fluorescent protein (GFP) is expressed under the control of the native promoter of *Mx1*, a type I IFN-induced gene that is transcribed but does not encode a functional protein in mice. The long half-life of GFP allows for sensitive and sustained detection of type I IFN-responsive cells. We confirmed in BMMs that GFP is strongly induced upon IFNβ (Figure S4A). On day 18 post-*Mtb* infection, we could detect type I IFN-responsive cells in lung lesions of *Sp140*^*–/–*^*Mx1*-GFP mice (Figures 4A, S4B), with much weaker reporter expression in B6.*Mx1*-GFP mice. As a key control, administration of anti-IFNAR1 antibody, previously shown to rescue the susceptibility of *Sp140*^−/−^ mice^35^, reduced *Mx1*-GFP reporter expression below the levels in B6, indicating that the *Mx1*-GFP reporter is specific and sensitive for type I IFN signaling *in vivo* in *Mtb*-infected mice (Figures 4A, S4B). We analyzed the lungs of B6 and *Sp140*^−/−^*Mx1*-GFP reporter mice at days 14-, 18- and 25 pi. While type I IFN signaling was detected in IMs as early as 14-days pi in both B6 and *Sp140*^−/−^ mice, the enhanced susceptibility of *Sp140*^−/−^ mice to *Mtb* only became apparent at day 25 pi (Figures 4B-C). Surprisingly, at day 14 and 18 pi, the frequency of type I IFN sensing IMs was similar across genotypes. However, by day 25 pi, the prevalence of GFP^+^ IMs decreased in B6 mice but was sustained in *Sp140*^*–/–*^ mice (Figure 4C). A similar sustained type I IFN response has recently been reported in C3HeB/FeJ susceptible mice^61^. The reporter allowed us to distinguish three levels of type I IFN signaling—none, weak and strong (Figure 4D). We defined weak expression as the level seen in B6 mice that is greater than background, whereas strong expression is the level seen in less than 5% of cells in infected B6 mice at day 18 pi. (but by many more cells in *Sp140*^*–/–*^ mice). At the early day 14 pi timepoint, the strength of IFN signaling in B6 and *Sp140*^−/−^ mice was similar (Figure 4E). However, by day 18 pi, elevated numbers of IMs responding strongly to type I IFN were present in *Sp140*^*–/–*^ versus B6 mice (Figure 4E). While the numbers of weak type I IFN signaling IMs dropped over time in both genotypes, cells exhibiting strong type I IFN signaling increased in number in *Sp140*^*–/–*^ mice at day 25 pi, but largely disappeared in B6 mice (Figure 4E). These observations demonstrate that at day 18 pi, a substantial subset of IMs from *Sp140*^*–/–*^ mice exhibit a stronger and more sustained type I IFN response. The presence of this subset precedes the appearance of elevated *Mtb* burdens in these mice.

**Figure 4.**
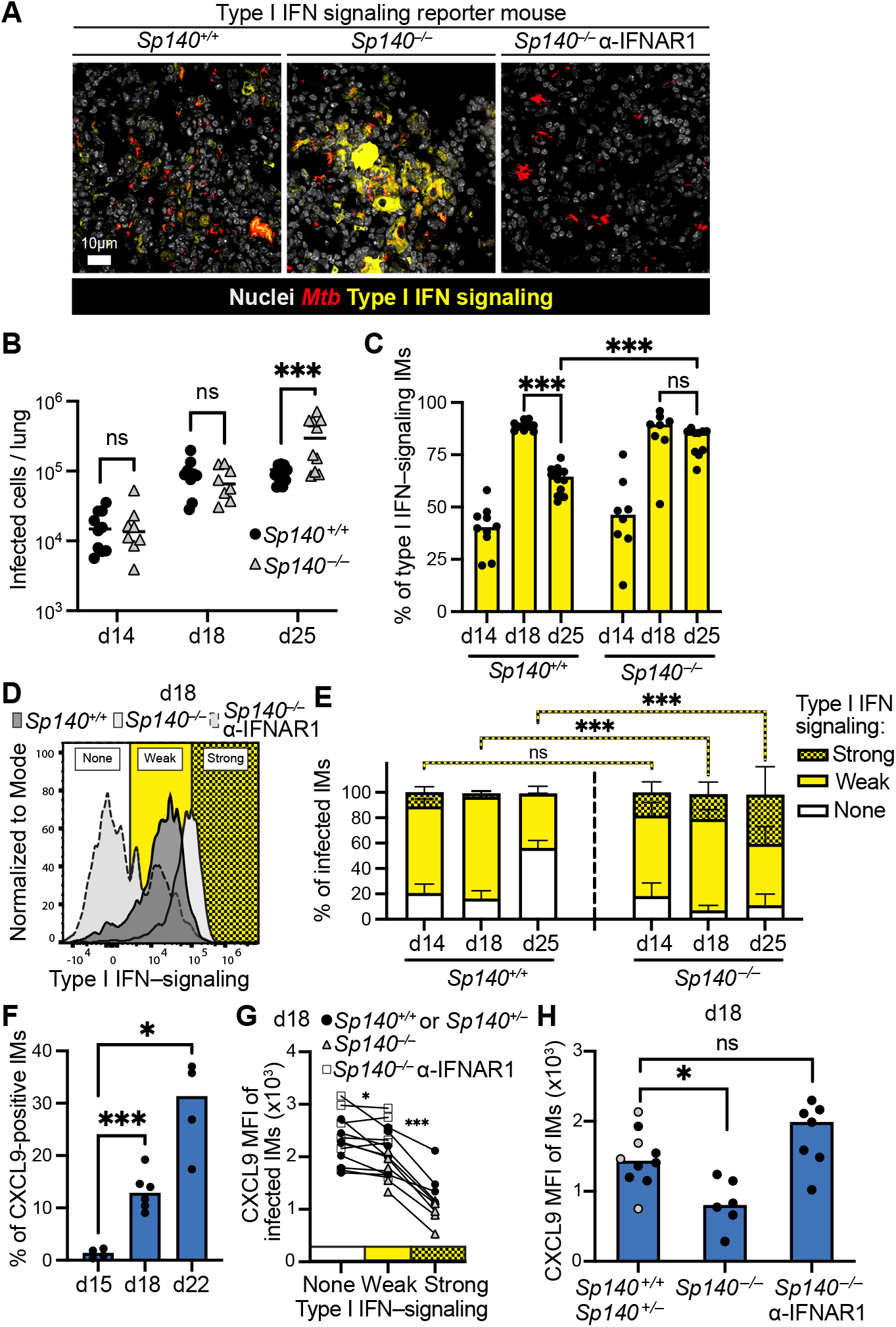
Sustained and strong type I IFN–signaling impairs the response to IFNγ and precedes susceptibility to *Mtb*. (**A**) Representative micrograph of lung tissue from *Mtb*-infected *Sp140*^*+/+*^ and *Sp140*^*–/–*^ Mx1-GFP reporter mice without or with administration of anti-IFNAR1 antibodies at day 18 pi. (**B**) Quantification of *Mtb*-infected cells and (**C**) Mx1-GFP-expression on IMs from lung tissue of *Sp140*^*+/+*^ and *Sp140*^*–/–*^ mice at day 14, 18 and 25 pi. (**D**) Representative flow cytometry histogram of Mx1-GFP expression of infected IMs from lung tissue of *Sp140*^*+/+*^ and *Sp140*^*–/–*^ mice without or with administration of anti-IFNAR1 antibodies at day 18 pi. “Weak” expression is defined as a level seen in B6 mice that is greater than background, whereas strong expression is the level seen in less than 5% of cells in infected B6 mice at day 18 pi (but by many more cells in *Sp140*^*–/–*^ mice). (**E**) Quantification of percent infected IMs with different levels of type I IFN signaling (as defined in (D)) from lung tissue of *Sp140*^*+/+*^ and *Sp140*^*–/–*^ mice at day 14, 18 and 25 pi. Data are represented as mean ± SD. (**F**) Percent IMs that are CXCL9^+^ from lung tissue of *Mtb*-infected B6 mice at day 15, 18 and 22 pi. (**G**) Comparison of CXCL9 MFI between none, weak and strong type I IFN signaling IMs (as defined in (D)) from lung tissue of *Sp140*^*+/+*^ *or Sp140*^*+/–*^ and *Sp140*^*–/–*^ mice without or with administration of anti-IFNAR1 antibodies at day 18 pi. (**H**) CXCL9 MFI of IMs from lung tissue of *Sp140*^*+/+*^ (black dots) or *Sp140*^*+/–*^ (gray dots) and *Sp140*^*–/–*^ mice without or with administration of anti-IFNAR1 antibodies at day 18 pi. (B-C, E-H) In each figure, data was combined from ≥ 2 biological replica with n ≥ 4 per group. Statistical significance was calculated in (B, C, E) with Two-way ANOVA and Sidak’s multiple comparison test, in (F, H) with Brown-Forsythe and Welch ANOVA test and Dunnett’s T3 multiple comparisons test, in (G) Mixed-effects analysis, with the Geisser-Greenhouse correction and with Sidak’s multiple comparison test with individual variances computed for each comparison. *p < 0.05, **p < 0.01, ***p < 0.001, ns = not significant.

In contrast to type I IFN signaling, the IFNγ response as assessed by CXCL9 expression was only apparent as early as 18-days pi (Figure 4F), which most likely coincides with the arrival of *Mtb*-specific T cells^62,63^. At this timepoint, we found that cells mounting a strong type I IFN response exhibited markedly decreased CXCL9 expression as compared to cells responding weakly or undetectably to type I IFNs (Figure 4G). The effect of strong type I IFN signaling on CXCL9 induction was evident in all *Sp140*^−/−^ mice (5 out of 5) and in the few B6 mice where we could detect some strong type I IFN signaling cells (4 out of 9, Figure 4G). However, because there were considerably more cells responding strongly to type I IFNs in *Sp140*^−/−^ mice, these mice exhibited an overall decrease in the levels of CXCL9 expression by IMs in the lung, as compared to B6 lungs (Figure 4H). At this timepoint (day 18 pi), CFU burdens were similar and CD4 T cells were equally recruited to *Mtb* within infected lesions of B6 and *Sp140*^−/−^ mice with or without anti-IFNAR1 antibody treatment (Figures S4C-E). However, at day ≥25 pi, CD4 T cells fail to localize to infected lesions in *Sp140*^−/−^ mice, whereas in B6 mice, CD4 T preferentially localize to the infected lesions (Figures S4F-G). Thus, type I IFN-dependent defects in IFNγ signaling preceded loss of bacterial control and dysfunctional localization of T cells.

Together, our results demonstrate that type I IFN signaling is apparent at early timepoints during *Mtb* infection, preceding the onset of the IFNγ response. In B6 mice, the type I IFN response is weak and transient; thus, responsiveness to IFNγ at later time-points is unimpaired, and bacterial restriction and T cell recruitment are intact. However, in *Sp140*^−/−^ mice, many cells respond strongly and persistently to type I IFN signaling, and these cells exhibit defective responsiveness to IFNγ, resulting in uncontrolled *Mtb* replication and impaired T cell recruitment, which might in turn further reduce the ability of IMs to restrict *Mtb* growth.

### RESIST promotes type I IFN responses and impairs IFNγ responses and *Mtb* control

Genetic or antibody-mediated suppression of IFNAR signaling rescues IFNγ signaling and susceptibility of *Sp140*^−/−^ mice to *Mtb*^35^ (Figures 2D, 4G-H). However, these conditions eliminate all type I IFN signaling and therefore do not specifically eliminate the sustained and strong type I IFN response that our data suggest is responsible for susceptibility to *Mtb*. We recently discovered RESIST (encoded by *Resist1* and *Resist2*) is a direct target of SP140 repression and is a crucial positive regulator that is required for enhanced (but not basal) IFNβ expression in *Sp140*^−/−^ mice^36^. To determine whether sustained and strong type I IFN signaling is the cause of type I IFN-driven susceptibility to *Mtb*, we tested if genetic loss of *Resist1* and *Resist2* (*Resist*^−/−^) rescues *Sp140*^*–/–*^ mice. Indeed, while *Resist*-deficiency had little impact in B6 mice (in which *Resist* is not normally expressed), *Sp140*^−/−^*Resist*^−/−^ mice were fully rescued for appropriate type I IFN signaling as compared to *Sp140*^*–/–*^ mice (Figure 5A). Loss of RESIST was also sufficient to restore the IFNγ response based on CXCL9 and NOS2 staining (Figures 5B, S5A). Consequently, RESIST-deficiency fully rescued the susceptibility of *Sp140*^*–/–*^ mice in terms of CFU burden (Figure 5C) and survival (Figure 5D). The rescue mediated by loss of *Resist* occurred without impairment of ‘normal’ IFNAR signaling. Thus, a normal (i.e., transient and modest) type I IFN response does not impair IFNγ responsiveness or *Mtb* control. Instead, our data demonstrate that strong and/or sustained type I IFN responses specifically mediate susceptibility to *Mtb* (Figure 5E).

**Figure 5.**
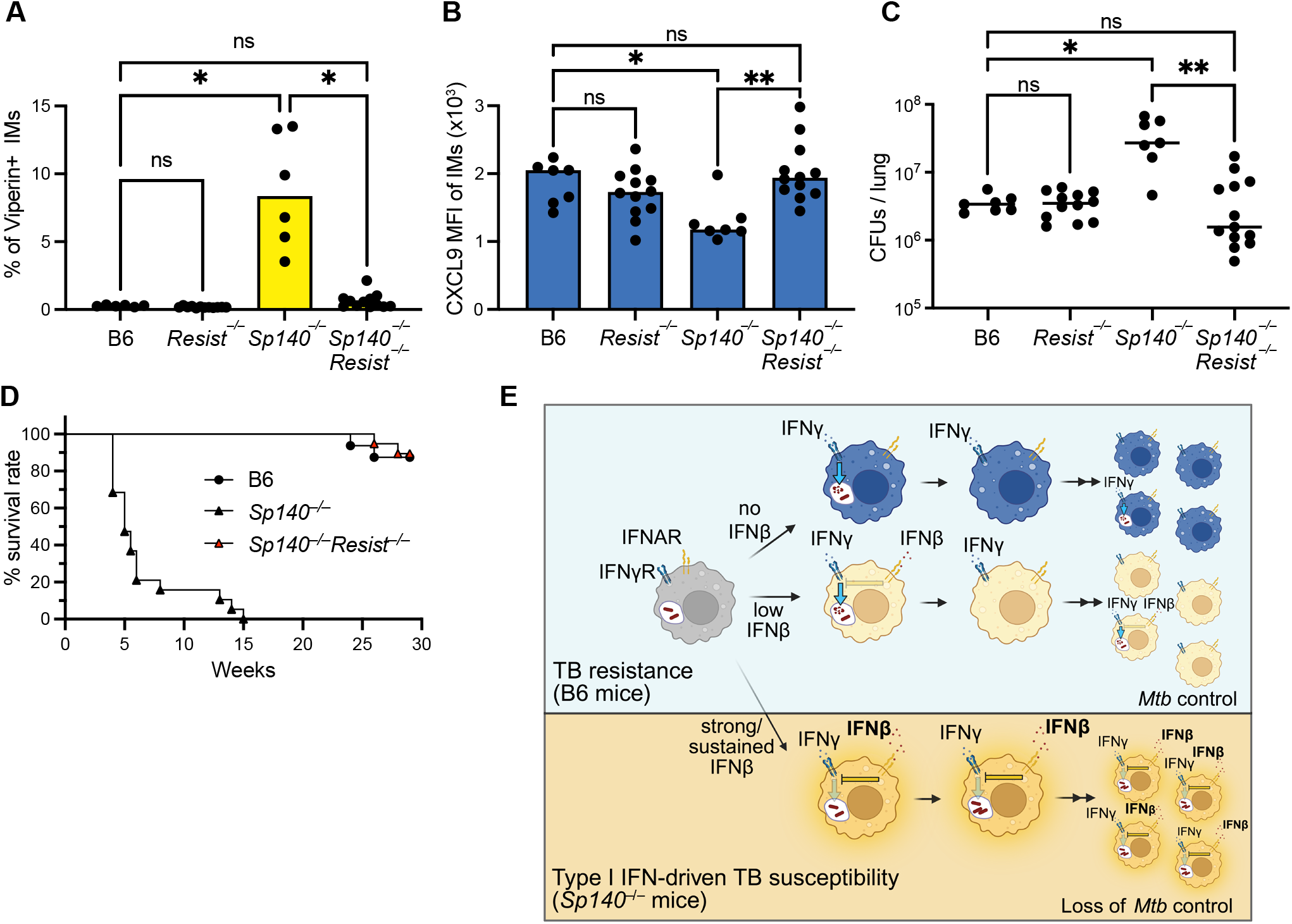
RESIST stabilizes and strengthens type I IFN responses, impairs IFNγ responses, and promotes susceptibility to *Mtb*. (**A**) Percent Viperin expression, (**B**) CXCL9 MFI of IMs and (**C**) CFUs from lung tissue of B6, *Resist*^*–/–*^, *Sp140*^*–/–*^ and *Sp140*^*–/–*^*Resist*^*–/–*^ mice at day 26-28 pi. (**D**) Survival experiment. (**E**) Model how sustained and strong type I IFN signaling impairs IFN**γ** responses and causes susceptibility to *Mtb*. Illustration created with BioRender. (A-D) Data was combined from ≥ 2 biological replicates with n ≥ 6 per group. Statistical analysis was calculated in (A-B) with Brown-Forsythe/Welch ANOVA test with Dunnett’s T3 multiple comparisons test and in (C) with Kruskal-Wallis test and Dunn’s multiple comparison test. *p < 0.05, **p < 0.01, ***p < 0.001, ns = not significant.

## Discussion

Type I IFNs consistently correlate with human TB disease progression^16–20^ and cause susceptibility to *Mtb* in numerous mouse models^21–30^. Previous studies have found that type I IFNs impair the protective IL-1 response via multiple mechanisms^29,31,64^. However, type I IFNs also promote TB susceptibility via IL-1-independent mechanisms^33,34,64^. Here, we provide evidence that sustained and strong (but not basal) type I IFN signaling cell intrinsically impairs IFNγ responsiveness in *Mtb*-infected macrophages in the lung. We further show that the impairment of IFNγ responsiveness is an early consequence of type I IFN signaling that renders IMs permissive for intracellular *Mtb* replication, initiating the loss of bacterial control and disease susceptibility. Previous *in vitro* studies have shown that type I IFNs can impair the induction of selected IFNγ-responsive genes^45–48^. However, many ISGs are induced by both type I IFNs and IFNγ, complicating efforts to clearly determine whether type I IFNs broadly suppress IFNγ responses. We show that IFNβ pre-exposure suppresses more than 90% of genes that are induced at least two-fold more strongly by IFNγ than by IFNβ. We also find that the impairment of IFNγ signaling persists well after active type I IFN signaling has subsided, requires the IFNAR-dependent signal transducers STAT2 and IRF9, and is conserved in both mouse and human primary macrophages.

In human *Mycobacterium leprae* infections, elevated type I IFN signatures correlate with impaired IFNγ responses and poor clinical outcomes^49^. For *Mtb*, we recently reported that impaired IFNγ responses correlate with elevated type I IFNs in *Sp140*^*–/–*^ mice^27^. However, a causal relationship has not previously been established. Based on our *in vitro* stimulation experiments and genetic evidence in mice, we identified CXCL9 as a robust marker of IFNγ responsiveness in IMs during *Mtb* infection (Figures 1, 2), consistent with its use in another recent study^65^. By comparing CXCL9 induction in *Mtb*-infected B6, *Sp140*^*–/–*^, and *Sp140*^*–/–*^*Ifnar1*^*–/–*^ mice at day 25-28 post infection, we found that type I IFN signaling causally impairs IFNγ responsiveness *in vivo* (Figure 2). However, at this time point (also used in prior studies^27^), bacterial burdens differ between genotypes, confounding our ability to distinguish a direct inhibitory effect of IFNAR signaling on IFNγ responses from indirect secondary effects of increased bacterial loads. To resolve this, we performed a time-course analysis (Figure 4), which revealed a clear temporal progression in which early type I IFN signaling precedes and suppresses the later-emerging IFNγ response, ultimately resulting in elevated bacterial burdens. We also analyzed mixed chimeric mice in which IFNAR-sufficient and -deficient cells respond in the same inflammatory environment. These experiments showed that IFNAR signaling cell-intrinsically represses IFNγ responsiveness and drives susceptibility to *Mtb* independent of bacterial burdens.

The mechanism by which type I IFNs impair IFNγ signaling has been studied extensively, but a single dominant mechanism has yet to emerge. It has been proposed that type I IFNs suppress IFNγ receptor expression^48,50^. Consistent with this, our RNA-seq analysis revealed a two-fold reduction in *Ifngr1* expression following IFNβ stimulation, and we previously reported reduced IFNγ receptor expression in SP140-deficient mice^27^. However, whether reduced receptor expression alone accounts for the observed impairment remains unclear. Additional, potentially redundant mechanisms are likely to contribute. For example, the shared signal transducer STAT1, which operates downstream of both IFNAR and IFNGR, may represent a limiting factor under conditions of sustained type I IFN signaling^66^. Moreover, IFNβ induces SOCS1, a well-established negative regulator of IFNγ signaling. Beyond these mechanisms, type I IFNs induce a broad antiviral response that includes ISGs known to suppress gene expression, such as PKR-mediated eIF2α phosphorylation, IFIT-dependent inhibition of translation initiation, and OAS–RNase L-mediated RNA degradation^67^. Thus, type I IFN-mediated suppression of IFNγ signaling is likely multifactorial, arising from the combined action of receptor-level regulation, shared signaling constraints, negative feedback pathways, and global repression of gene expression.

A major unresolved question has been why type I IFNs promote TB pathogenesis in certain mouse strains (e.g., C3HeB/ FeJ^29,68^ or 129^69^), but not in B6 mice. This has been puzzling because B6 mice are fully capable of mounting a robust type I IFN response^70^, and indeed, are generally highly resistant to viral infections^71–73^. Previous data have shown that type I IFNs are expressed at higher levels in susceptible mouse strains during *Mtb* infection, but since *Mtb* itself induces type I IFNs^40,74^, it has been difficult to determine if mice are susceptible due to higher interferon levels, or if higher bacterial burdens in susceptible mice are what cause the increased interferon responses. Prior experiments have shown that deleting *Ifnar* in susceptible mice^29,69^ restores resistance, but these experiments do not address whether elevated IFN levels drive susceptibility, because *Ifnar* deletion eliminates all type I IFN signaling and does not selectively eliminate the elevated levels while maintaining basal interferon responses. To gain insight into the quantitative dynamics by which type I IFN signaling impairs IFNγ responses and promotes susceptibility to *Mtb*, we crossed *Sp140*^*–/–*^ mice to mice harboring the sensitive *Mx1*-GFP reporter, in which type I IFN signaling induces expression of a stabilized GFP^60^. With these mice, we show that IFNγ responsiveness is not determined simply by the presence or absence of type I IFN signaling but instead depends critically on the magnitude and/or duration of the response (Figure 4). Specifically, strong and/or sustained type I IFN signaling suppresses IFNγ signaling and promotes susceptibility to *Mtb*. This was not solely observed in *Sp140*^*–/–*^, but also to a limited extend in B6 mice, in line with a recent report using *Mtb* strains that induced early detrimental type I IFN responses in B6 mice^61^. Our time course analysis shows that the enhanced type I IFN response in *Sp140*^−/−^ mice versus wild-type mice occurs prior to the divergence in bacterial burdens, and is thus not a consequence of elevated bacterial burdens, but is instead an important driver of susceptibility to *Mtb*. This conclusion is supported not only by data from the *Mx1*-GFP reporter mice, but also from independent genetic experiments (Figure 5) in which we deleted the *Resist* locus that we previously showed encodes a positive regulator of type I IFN production that is specifically de-repressed in *Sp140*^−/−^ mice^36^. *Resist* deletion had no detectable effect in the B6 background (in which it is not normally expressed) and did not affect basal type I IFN levels. However, in *Sp140*^*–/–*^ mice, RESIST deficiency restored an appropriately transient type I IFN response, thereby permitting IFNγ-dependent restriction of *Mtb* comparable to that observed in B6 mice.

Our findings emphasize the key point that IFNγ production does not necessarily translate into effective IFNγ signaling. Instead, IFNγ responsiveness is shaped by the surrounding cytokine environment, particularly by the presence of type I IFNs. Accordingly, assessment of IFNγ activity requires measurement of downstream responses, rather than IFNγ levels alone. Here, we provide robust markers to reliably quantify IFNγ responsiveness and offer a clear example in the context of TB of how this distinction is critical. Together, our findings may help explain why induction of IFNγ or IFNγ-producing CD4^+^ T cells^11–15^ is not always sufficient to confer protection against *Mtb* or other bacterial pathogens in which type I IFN-driven susceptibility is observed. We also reveal potential strategies for enhancing IFNγ-mediated immunity that could lead to more effective host directed therapies.

## Materials and Methods

### Mice

Mice were maintained in accordance with the regulatory standards of the University of California Berkeley Institutional Animal Care and Use Committee under specific pathogen-free conditions. In all experiments mice were age- and sex-matched and were 8-18 weeks old at the start of the infections. Every experiment included female and male mice. Mouse lines used are C57BL/6J (B6), B6.129S2-Ifnar1^tm1Agt^/Mmjax (*Ifnar1*^*–/–*^), B6.129S7-Ifngr1tm1Agt/J (*Ifngr1*^*–/–*^), C57BL/6J-Ptprcem6Lutzy/J (CD45.1), B6.Cg-Mx1tm1.1Agsa/J (*Mx1*^*g fp*^), B6(Cg)-Ifnar-1tm1.1Ees/J (*Ifnar1*^*fl/fl*^) and C57BL/6N-Ifngr1tm1.1Rds/J (*Ifngr1*^*fl/ fl*^) that were purchased from Jackson Laboratories. *Sp140*^*–/–*^ previously made^35^ was crossed in house to *Ifnar1*^*–/–*^, CD45.1 and *Mx1*^*g fp*^ to generate *Sp140*^*–/–*^ *Ifnar1*^*–/–*^, *Sp140*^*–/–*^CD45.1 and *Sp140*^*–/–*^*Mx1*^*g fp*^, respectively. B6-Fcgr1tm2Ciphe (CD64^Cre^) was previously described^75^. *Ifnar1*^*fl/fl*^ and *Ifngr1*^*fl/fl*^ were crossed to *Sp140*^*–/–*^ and CD-64^Cre^ to generate *Sp140*^*–/–*^*Ifnar1*^*fl/fl*^ CD64^Cre^ and *Ifngr1*^*fl/f*^ CD64^Cre^. *Sp140*^*–/–*^*Resist*^*–/–*^ were previously described ^36^ and backcrossed in house to B6 to generate *Resist*^*–/–*^. *Sp140*^*–/–*^*Irf9*^*–/–*^ and *Sp140*^*–/–*^ *Stat2*^*–/–*^ mice were generated in this study as described below.

### Mouse bone marrow macrophages (BMMs)

Bones (femurs and tibias) from B6 or *Ifnar1*^*–/–*^ mice were harvested and sterilized in 70% EtOH. Bone marrow was isolated by flushing the bones with ice-cold BMM media consisting of DMEM supplemented with 10% fetal bovine serum (FBS), 10% MCSF (generated from 3T3 cells), GlutMax, 10mM HEPES and Pen-Strep (Thermo). Isolated bone marrow was filtered through a 70µm cell strainer, centrifuged at 4°C for 5 min, 600G, resuspended in BMM media and seeded into 6-8 15cm non-treated petri dishes. Cells were incubated for 7 days to differentiate into BMMs with adding 50% BMM media on day 3 before transferring cells into the appropriate multi-well format using a cell scraper. After 2 days of resting, BMMs were exposed to cytokines according to the experimental setup.

### Human macrophages

THP-1 cells (ATCC) were maintained in RPMI including 10% FBS, GlutMax and Pen-Strep (complete RPMI). THP-1 were differentiated by adding 100 ng/mL phorbol myristate acetate (PMA, Invivogen, tlrl-pma) for 48h followed by 36h rest before used for exposure assays. Cryopreserved negatively selected primary human monocytes were purchased from AllCells and differentiated for 6 days in complete RPMI supplemented with 50 ng/ml human M-CSF (PeproTech, 300-25). Cells were lifted with trypsin and transferred into a 96-well format for exposures.

### Generation of *Sp140*^*–/–*^*Irf9*^*–/–*^ and *Sp140*^*–/–*^*Stat2*^*–/–*^ mice

*Sp140*^*–/–*^*Irf9*^*–/–*^ and *Sp140*^*–/–*^*Stat2*^*–/–*^ mice were generated by electroporation of Sp140–/– zygotes with Cas9 and sgRNA UACG-CUGCACCCGAAAGCUG and AGUGGUCCCACUGGUU-CAGU, respectively. Founders were genotyped and backcrossed to *Sp140*^*–/–*^ mice, and progeny with matching alleles were further bred. Genotyping was performed by sequencing of PCR product with the primer pairs CAGGGGTTTGCAAGTTGTTG, AGACATGGTTGGTTCTACTTTCT for *Irf9* and GGCT-CATCTGATTTCAGGCC, CCTCTCAGGTGACACACAAC for *Stat2*. Established knock-out mouse lines had a 20, 17 base pair deletion in *Irf9* exon 3, *Stat2* exon 3, respectively.

### Mouse *Mtb* infections, *in vivo* antibody mediated blockage, tissue processing for CFU and flow cytometry analysis

*Mtb* Erdman strains including the ones constitutively expressing either mWasabi or mCherry have previously been described^27^. Inoculum was prepared from a frozen stock and diluted in 9ml sterile PBS at an of ca. OD of 0.002. Mice were inserted into an aerosolizer device (Glas-Col, Terre Haute, IN) and infected through the aerosol route at a low dose of ca. 20-100 CFUs.

Infectious dose was verified from 2-3 mice at day 1 post infection. For IFNAR1 blockage *in vivo* 500ug anti-IFNAR1 antibody (bioXcell, MAR1-5A3) per mouse was injected intraperitoneal every other day starting at day 7 pi. Mice were sacrificed at indicated time points post infections and the complete lungs were harvested into a GentleMACS C tube (Miltenyi Biotec) with 2ml digestion media consisting of 2ml RPMI media (Gibco) with 30ug/ml DNase I (Roche), 70ug/ml Liberase TM (Roche) and Brefeldin A (BioLegend). Lungs were cut into large pieces using the program lung_01 on GentleMACS device (Miltenyi Biotec) before incubating at 37°qC for 30 minutes. Lungs were homogenized with program Lung_2 on the GentleMacs device and digestions was stopped by adding 2ml of PBS including 20% Newborn Calf Serum (Thermo Fisher Scientific). Lung homogenate was filter through a 70µm SmartStrainers (Miltenyi Biotec) into 15ml falcon tube. From this single cell suspension 100ul was saved for CFU plating assay, whereas the rest centrifuged at 1,600 rpm for 8 minutes at 4°C. Cell pellet was resuspended in FACS buffer and used for flow cytometry analysis.

### CFU plating

7H11 plates supplemented with 10% BD BBL™ Middlebrook OADC Enrichment (Fisher) and 0.5% glycerol were prepared ahead of time and stored at 4°C. Lung homogenates were serial diluted in PBS and 50ul of appropriate dilutions were plated. After 3 weeks incubation at 37°C, CFUs were enumerated to back calculate CFUs per lung.

### Flow cytometry of lung homogenate

The following fluorophore-coupled antibodies were used accordingly together with fixable viability dye (Ghost Dye™ Violet 780; Tonbo Biosciences), TruStain FcX PLUS (S17011E, BioLegend), Super Bright Complete Staining Buffer (Thermo Fisher Scientific) and True-Stain Monocyte Blocker (BioLegend) to prepare a Master Mix and stain the single cells suspension from above: BV421-coupled MHCII (M5/114.15.2, BioLegend), BV480-coupled B220 (RA3-6B2, BD Biosciences), BV480-coupled CD90.2 (53-2.1, BD Biosciences), BV605-coupled CD64 (X54-5/7.1, BioLegend), BV711-coupled CD11b (M1/70, BioLegend), BV785-coupled Ly6C (HK1.4, BioLegend), PE-Cy7-coupled MerTK (DS5MMER, Thermo Fisher Scientific), APC-R700-coupled Siglec F (E50-2440, BD Biosciences), BUV496-coupled CD45 (30-F11, BD Biosciences), BUV563-coupled Ly6G (1A8, BD Biosciences), BUV737-coupled CD11c (HL3, BD Biosciences), BV421-coupled CD45.1 (A20, BioLegend), BUV496-coupled CD45.2 (104, BD Biosciences). Staining was performed at room temperature for >30 minutes before washing the cells three times with FACS buffer. Stained cells were fixed with Cytofix/Cytoperm (BD Biosciences) for >30 minutes at room temperature before retrieved from BSL3 facility. Fixed cells were permeabilized by washing cells four times with Permeabilization Buffer (Invitrogen) before performing intracellular staining using PE-coupled Viperin (MaP.VIP, BD Biosciences), AF647-coupled CXCL9 (MIG-2F5.5, BioLegend) and BUV395-coupled NOS2 (CXNFT, Invitrogen). Cells were run on an Aurora (Cytek) flow cytometer and analyzed with Flowjo version 10 (BD Biosciences).

### Confocal Microscopy for imaging and image analysis

For microscopy the middle lobe was harvested and directly fixed with Cytofix/Cytoperm (BD Biosciences) diluted in PBS (1:2) for >24h at 4°C before retrieving from BSL3 laboratory. Fixed lung tissues were washed with PBS and dehydrated for >12h at 4°C in PBS containing 20% sucrose. Lung tissues were embedded in O.C.T. (Tissue-Tek) and stored at −80°C. 10µm sections were prepared using a Leica CM3050S Cryotome and mounted on Superfrost Plus Microscope Slides (Fisher). Sections were rehydrated with PBS, permeabilized with PBS containing 0.5% Tx-100 and blocked with 10% Normal Goat Serum (Vector Laboratories) before staining with DAPI (Sigma) and fluorophore-coupled antibodies. Stained tissue was covered with a cover slip and VectaShield HardSet (Vector Laboratories). A Zeiss LSM710 confocal microscope and FIJI software was used for image analysis. Shortest distance analysis was performed with a FIJI plugin for distance analysis called DiAna^76^.

### Confocal microscopy and image processing for histo-cytometry analysis

Confocal Microscopy was performed using a Zeiss LSM 880 laser scanning confocal microscope (Zeiss) equipped with two photomultiplier detectors, a 34-channel GaASP spectral detector system, and a 2-channel AiryScan detector as well as 405, 458, 488, 514, 561, 594, and 633 lasers. Stained 20μm paraformaldehyde fixed lung sections from *Mtb*-mCherry infected mice were inspected with a 5× air objective to find representative lesions and distal sites and then imaged using a 63× oil immersion objective lens with a numerical aperture of 1.4. For each infected lung, one 12-15 x 12-15 tiled Mtb-heavy lesion image and one 4×4 tiled distal site image was taken consisting of 20μm z-stacks with a 1.5µm step size. Additionally, the Zeiss LSM 880 microscope was used to image single color-stained Ultracomp eBeads Plus (Thermo Fisher Scientific) for generating a compensation matrix. Image analysis was performed using Chrysalis software.^77^ Briefly, a compensation matrix was generated by automatic image-based spectral measurements on single col-or-stained controls in ImageJ by using Generate Compensation Matrix script. This compensation matrix was used to perform linear unmixing on three-dimensional images with Chrysalis. Chrysalis was also used for further image processing, including rescaling data and generating new channels by performing mathematical operations using existing channels. For histo-cytometry analysis, Imaris 9.9.1 (Bitplane) was used for surface creation to digitally identify cells in images based on protein expression.^78^ Total number, surface volume, and mean fluorescence intensity of each cell type within a given surface were calculated, along with total volume of each z-stacked section. Statistics for identified cells were exported from Imaris and then imported into FlowJo version 10 (BD Biosciences) for quantitative image analysis.

### Bone marrow chimeras

From donor mice, bones (femurs and tibias) were harvested, sterilized in 70% EtOH and bone marrow was isolated by flushing the bones with cold PBS using a syringe and filtering through a 70µm cell strainer. Cells were once washed and resuspended in PBS for injection. For mixed bone marrow chimeras, bone marrow cells from different genotypes were mixed in a 1:1 ratio. Recipient mice were lethally irradiated two times within 12-20 hours with a Precision X-Rad320 X833 Ray irradiator (North Branford, CT). Mice received 2-5 Mio bone marrow cells in 200ul PBS by retro-orbital injection. Mice were housed for >8 weeks to allow hematopoietic reconstitution prior *Mtb* infection.

### Cytokine exposures of cells for flow cytometry analysis

For exposures, cytokines expressed in either HEK293 or CHO cells were used. Specifically, if not other stated in figures or figure legends, 2ng/ml mouse IFNβ (581304, BioLegend), 50ng/ ml mouse IFNγ (ab259378, Abcam), 10ng/ml mouse TNF (ab259411, Abcam), 2ng/ml human IFNβ (300-02BC, Thermo), 50ng/ml human IFNγ (ab259377, Abcam). TLR agonist used were 50ng/ml Pam3CSK4 (Invivogen, tlrl-pms) and 10ng/ml LPS (Invivogen, tlrl-3pelps).

### Flow cytometry of cytokine exposed macrophages

Brefeldin A was added to the BMM media for the last 4 hour of cytokine exposure. BMM media was removed, and BMMs were incubated for 20min with pre-warmed PBS with 4mM EDTA. Detached BMMs were transferred and washed once in PBS prior staining with fixable viability dye (Ghost Dye™ Violet 780; Tonbo Biosciences). After 30min incubation at room temperature, cells were washed three times with PBS and fixed with IC Fixation Buffer (eBioscience) for 15 minutes at room temperature. Cells were washed and fixed four times with Permeabilization Buffer (Invitrogen) before performing intracellular staining using PE-coupled Viperin (MaP.VIP, BD Biosciences), AF647-coupled CXCL9 (MIG-2F5.5, BioLegend) and AF488-coupled NOS2 (CXNFT, eBiosciences). Cells were run on a Fortessa (BD Biosciences) flow cytometer and analyzed with Flowjo version 10 (BD Biosciences). All the analysis were performed on viable cells only.

### Cas9-ribonucleoprotein (RNP) mediated gene disruption in BMMs

Gene disruption in BMMs was performed with Cas9-ribonucleoprotein (RNP) electroporation at day 5 post seeding as described previously^79^. In brief, Cas9 2 NLS nuclease (Synthego) was pre-incubated with gRNAs (Synthego, sgRNA EZ kits) and Alt-R Cas9 Electroporation Enhancer (IDT, 1075916) for >20min at room temperature to form Cas9-RNPs. BMMs were lifted using a cell scraper, washed in PBS and resuspended in Lonza P3 buffer (Lonza, V4XP-3032) including Supplement 1 according to manufacturer’s protocol before combining with Cas9-RNPs. Cells were electroporated with the Lonza 4D-Nucleofector Core Unit (AAF-1002B) using the program CM-137. Electroporated BMMs were recovered in BMM media and 2Mio cells were plated per 10cm non-treated petri dish and incubated at 37°. 50% fresh media was added two days post electroporation and after 4 days of recovery edited BMMs were transferred into the appropriate multi-well format for exposure assays. The following gRNA sequences were used: Ifnar2: CAGACGGU-GUGAUAGUCUCU & CAAAGACGAAAAUCUGACGA, Stat2: AGUGGUCCCACUGGUUCAGU, Irf9: UACGCUG-CACCCGAAAGCUG & GUUGUAAACCACUCAGACAG.

### Bulk RNA-seq sample preparation and analysis

For exposures 10ng/ml mouse IFNβ (581304, BioLegend), 10ng/ ml mouse IFNγ (ab259378, Abcam) was used. Upon cytokine exposure, BMMs were lysed with TRK lysis buffer (Omega Bio-Tek) including 2-mercaptoethanol (Thermo Fisher Scientific).

Total RNA isolation was performed using the E.Z.N.A Total RNA Kit I (Omega Bio-Tek) with a DNase treatment on-column (Qiagen, 79254). The library preparation, sequencing, and read alignment to the mouse genome was performed by Azenta Life Sciences. Raw counts were used as input for analysis with DESeq2 ^80^.

### RT-qPCR analysis

Total RNA was isolated the same way as for bulk RNAseq described above. cDNA was reverse transcribed from RNA with Superscript III Reverse Transcriptase (Invitrogen, 18080093) and oligo dT18 (NEB, S1316S) in the presence of RNase inhibitors. Diluted cDNA was assessed by RT-qPCR using the Power SYBR Green PCR Master Mix (Thermo Fisher Scientific, 43-676-59) in technical duplicates. PrimeTime qPCR Primers (IDT) were used: Cxcl9 Mm.PT.58.5726745, Rsad2: Mm.PT.58.11280480, Actin: Mm.PT.39a.22214843.g.

### Statistical Analysis

Statistical tests to determine statistical significance were performed using Prism (GraphPad) software and are indicated in the figure legends. *p < 0.05, **p < 0.01, ***p < 0.001, ns = not significant.

## Acknowledgments

We thank members of the Vance, Barton, Stanley, and Cox laboratories for helpful discussions, the UC Berkeley Cancer Research Laboratory Flow Cytometry facility for assistance with flow cytometry, the UC Berkeley Biological Imaging Facility for assistance with microscopy, and the UC Berkeley Office of Laboratory Animal Care for housing the mice. The model figure was created with BioRender.com.

## Funding

SAF was supported by an EMBO Postdoctoral Fellowship (ALTF 617-2021) and a Postdoc Mobility-Fellowship from the Swiss National Science Foundation (P500PB_206801). R.E.V. is an HHMI Investigator and is supported by NIH grants AI075039, AI066302, and AI155634.

## Author contributions

Conceptualization: S.A.F. and R.E.V.; investigation: S.A.F., B.P., R.C., M.R.F., O.V.L., E.A.T., E.B., and D.I.K.; data analysis: S.A.F. and B.P.; methodology: S.A.F., K.C.W., J.J.R. and D.I.K.; resources: H.D. and A.Y.L.; supervision: S.A.F. and R.E.V. Funding acquisition: S.A.F. and R.E.V. writing: S.A.F. and R.E.V.

## Competing interests

R.E.V. consults for and is on the Scientific Advisory boards of X-biotix Therapeutics, Ditto Biosciences, and Remedy Plan, Inc.

---

**Figure S1.**
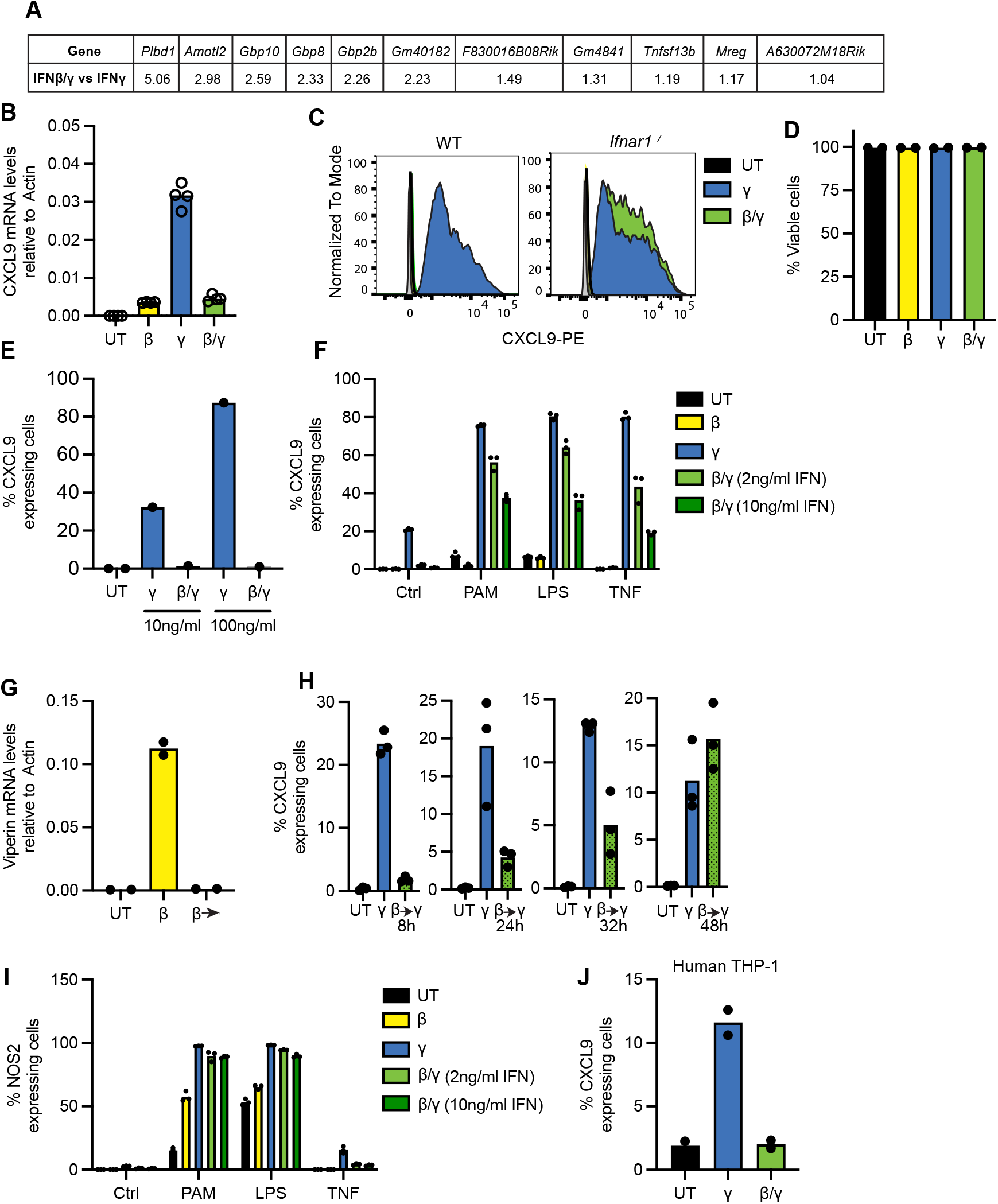
Supplementary to Figure 1. (**A**) Table of the IFN**γ** specific genes that are not inhibited by type I IFN signaling. (**B**) Quantification of CXCL9 mRNA levels by RT-qPCR from BMMs exposed to IFN**γ** with or without IFNβ-preexposure. UT = Untreated (**C**) Representative flow cytometry histogram of CXCL9 from WT and *Ifnar1*^*–/–*^ BMMs exposed to IFN**γ** with or without IFNβ-preexposure. (**D**) Quantification of viable BMMs upon exposure to cytokines. (**E**) Quantification of CXCL9 protein expressing BMMs from WT mice exposed to 10 and 100ng/ml IFN**γ** with or without IFNβ-preexposure. (**F**) Quantification of CXCL9 protein expressing BMMs from WT mice upon exposure to IFN**γ**, IFNβ or IFNβ/**γ** in the absence or presence of tumor necrosis factor (TNF) and Toll-like receptor agonists Pam3CSK4 (PAM) or Lipopolysaccharide (LPS). (**G**) Quantification of *Rsad2* mRNA (encoding Viperin) levels by RT-qPCR of WT BMMs directly upon IFN**γ** exposure or after removing it for 8h. (**H**) Quantification of CXCL9 protein expressing BMMs from WT mice exposed to IFN**γ** at various time post transient overnight IFNβ-exposure. (**I**) Quantification of NOS2 protein expressing BMMs from WT mice upon exposure to IFN**γ**, IFNβ or IFNβ/**γ** in the absence or presence of TNF and Toll-like receptor agonists PAM or LPS. (**J**) Quantification of CXCL9 protein expressing human THP-1 macrophages exposed to IFN**γ** with or without IFNβ-preexposure. (B, D-J) Representative data from ≥ 3 biological replica with (B, D, F-J) n ≥ 2 and with (E) n = 1 per condition.

**Figure S2.**
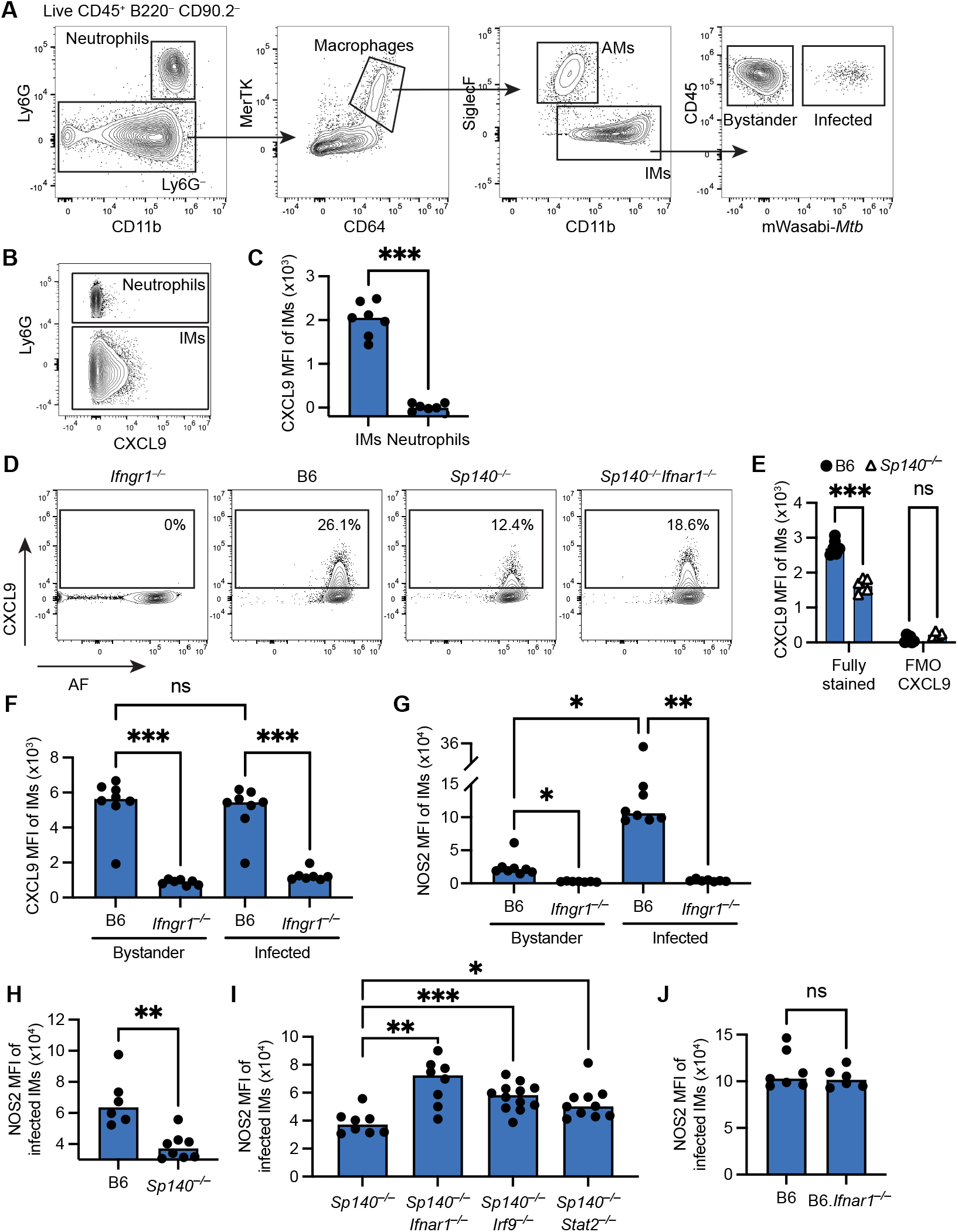
Supplementary to Figure 2. **(A)** Representative flow cytometry plots of a *Mtb*-mWasabi infected B6 mouse gated on live CD45^+^ B220^−^ CD90.2^−^ cells to identify interstitial macrophages (CD64^+^ MerTK high SiglecF^−^), which can be grouped into bystander (mWasabi^−^) and infected (mWasabi^+^) cells. **(B)**Representative flow cytometry plots of CXCL9 expression by IMs and neutrophils at day 28-29 pi and in (**C**) corresponding quantification. (**D**) Representative flow cytometry plots of CXCL9 vs autofluorescence (AF) in indicated mouse strains at day 25-26 pi. (**E**) Quantification of CXCL9 MFI of IMs in fully stained and fully stained minus CXCL9 (FMO CXCL9) samples from *Mtb*-infected B6 and *Sp140*^*–/–*^ mice at day 25 pi. (**F**) MFI of CXCL9 and (**G**) MFI of NOS2 in IMs from *Mtb*-infected B6 and *Ifngr1*^*–/–*^ mice at day 25 pi grouped in bystander and infected IMs. (**H**) MFI of NOS2 staining in IMs from *Mtb*-infected B6 and *Sp140*^*–/–*^ mice at day 25 pi. (**I**) MFI of NOS2 staining in IMs from *Mtb*-infected *Sp140*^*–/–*^, *Sp140*^*–/–*^*Ifnar1*^*–/–*^, *Sp140*^*–/–*^*Irf9*^*–/–*^ and *Sp140*^*–/–*^*Stat2*^*–/–*^ mice at day 28-29 pi. (**J**) MFI of NOS2 staining in IMs from *Mtb*-infected B6 and B6.*Ifnar1*^*–/–*^ mice at day 25-26 pi. (C, E, F-J) In each figure, data was combined from ≥ 2 biological replica with n ≥ 6 per group. Statistical significance was calculated in (E-G) with Two-way ANOVA and Sidak’s multiple comparison test, in (I) with Brown-Forsythe and Welch ANOVA test and Dunnett’s T3 multiple comparisons test, in (C, H, J) with Welch’s t-test. *p < 0.05, **p < 0.01, ***p < 0.001, ns = not significant.

**Figure S3.**
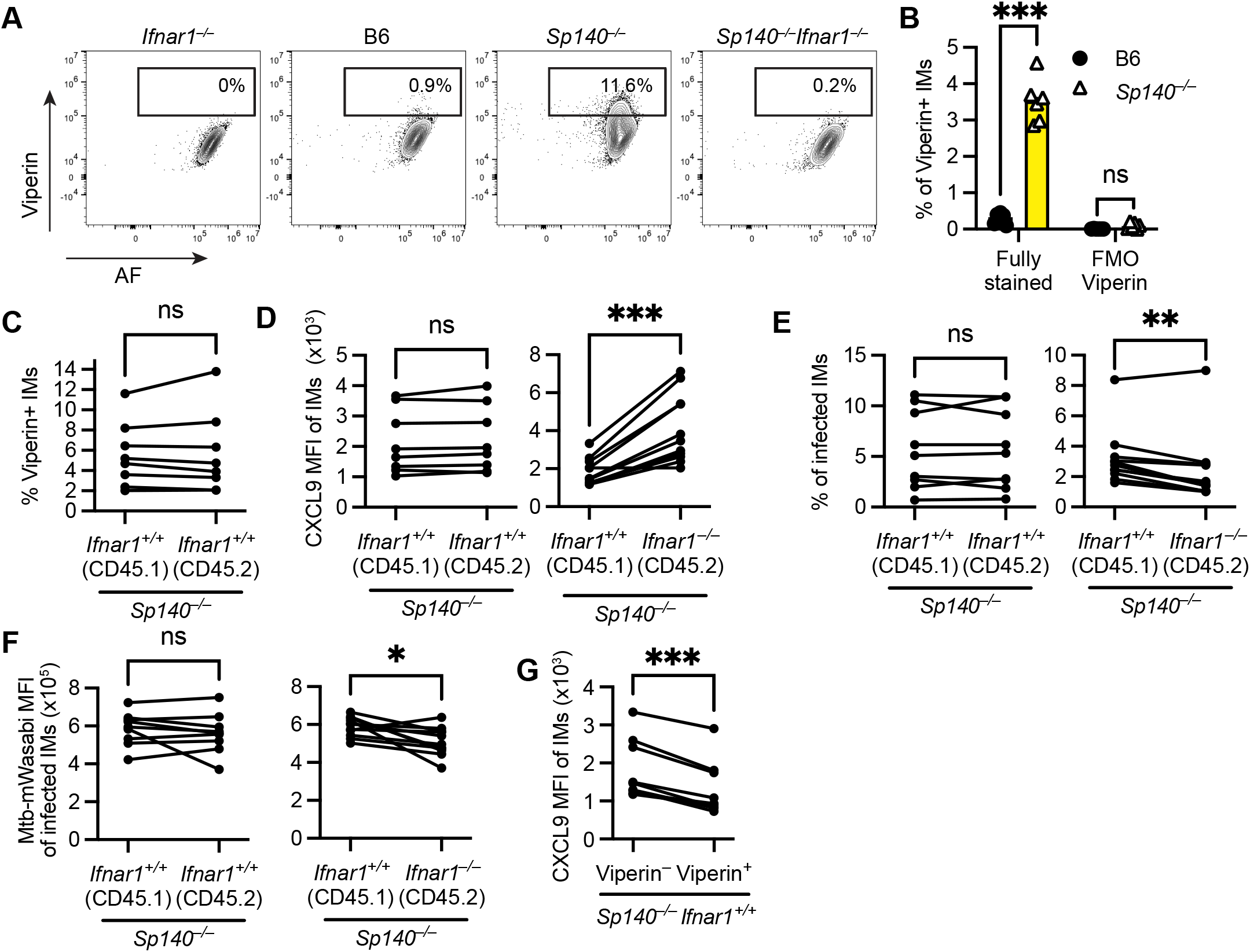
Supplementary to Figure 3. (**A**) Representative flow cytometry plots of Viperin vs autofluorescence (AF) in indicated mouse strains at day 25-26pi. (**B**) Quantification of Viperin MFI of IMs in fully stained and fully stained minus Viperin (FMO Viperin) samples from *Mtb*-infected B6 and *Sp140*^*–/–*^ mice at day 25 pi. (**C-F**) Analysis of *Mtb*-infected mixed *Sp140*^*–/–*^ bone marrow chimeras containing CD45.1- and CD45.2-positive cells, either both IFNAR-proficient (control) or IFNAR-proficient and IFNAR-deficient, respectively, at day 25-26 pi. Comparison in terms of (**C**) Viperin positivity, (**D**) CXCL9 MFI, (**E**) percent infected IMs and (**F**) *Mtb*-mWasabi MFI. (**G**) CXCL9 MFI analysis of Viperin-negative and -positive IMs from *Sp140*^*–/–*^*Ifnar1*^*+/+*^ subpopulation in figure 3J. Data in (B) from one biological replica with n = 5 per group and in (C-G) from ≥ 2 biological replica with n ≥ 6 per group. Statistical significance was calculated in (B) with Two-way ANOVA and Sidak’s multiple comparison test, in (C-G) with paired t-test. *p < 0.05, **p < 0.01, ***p < 0.001, ns = not significant.

**Figure S4.**
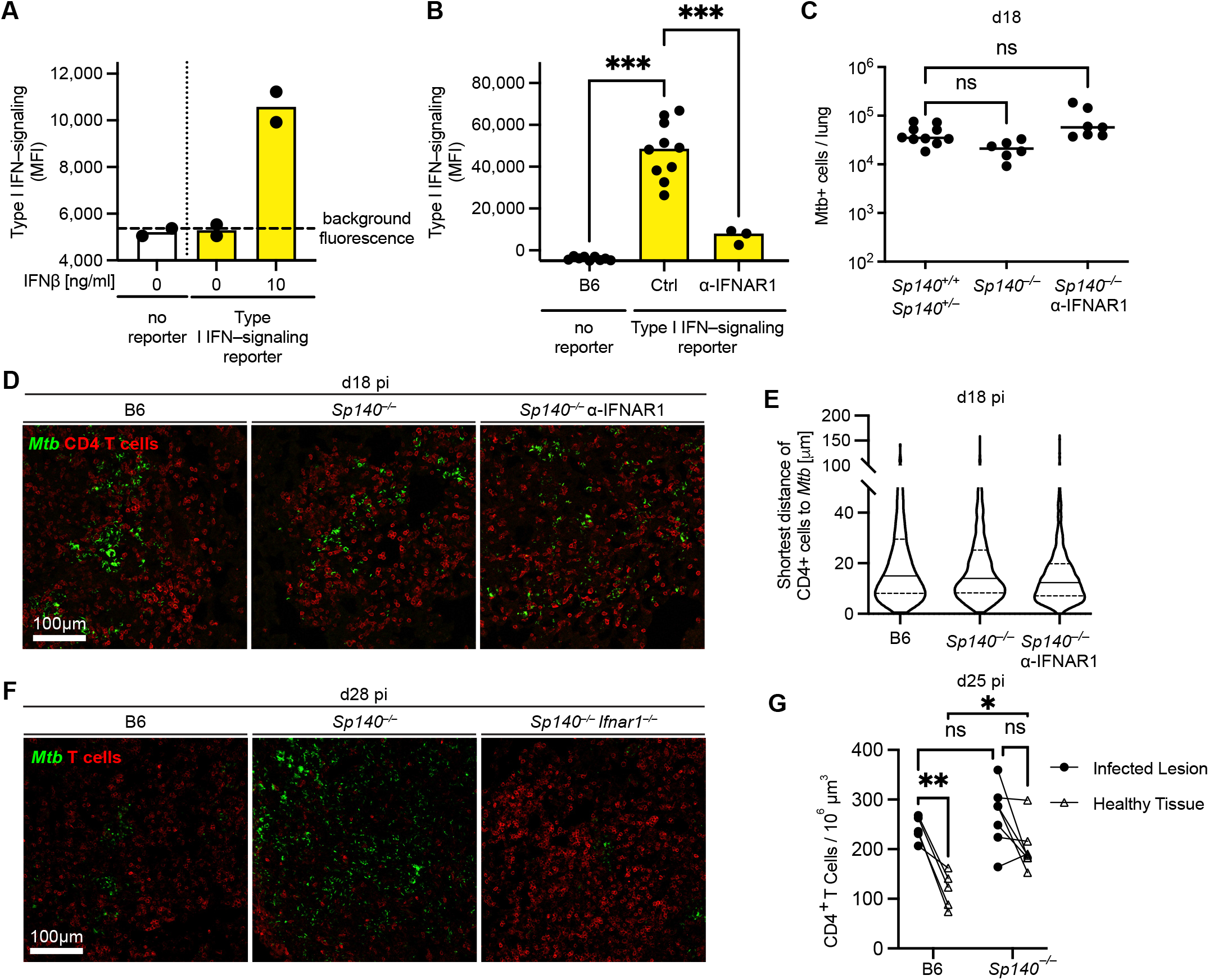
Supplementary to Figure 4. (**A**) Quantification of Mx1-expression of BMMs established from type I IFN-signaling reporter mouse line (*Mx1*^*gfp*^) upon exposure to IFNβ compared to background fluorescence. (**B**) MFI of Mx1-GFP expression in IMs from *Mtb*-infected *Mx1*^*gfp*^ mice without or with administration of anti-IFNAR1 antibodies. (**C**) Quantification of *Mtb*-mCherry-positive cells from *Mtb*-infected *Sp140*^*+/+*^, *Sp140*^*+/–*^, and *Sp140*^*–/–*^ mice without or with administration of anti-IFNAR1 antibodies at day 18 pi. (**D**) Representative micrograph with CD4^+^ T cells of infected lung lesions from *Mtb*-infected B6, *Sp140*^*–/–*^ and *Sp140*^*–/–*^ treated with anti-IFNAR1 antibodies at day 18 pi. (**E**) Shortes distance quantification of CD4^+^ T cells to *Mtb* within infected lesions at day 18 pi. (**F**) Representative micrograph with T cells of lung lesions from *Mtb*-infected B6, *Sp140*^*–/–*^ and *Sp140*^*–/–*^ treated with anti-IFNAR1 antibodies at day 28 pi. (**G**) Quantification of CD4^+^ T cells per 10^6^ µm^3^ in “Infected Lesions” and “Healthy Tissue” from B6 and *Sp140*^*–/–*^ *Mtb*-infected mouse lungs at day 25 pi. Data in (A) from one biological replica with n = 2 per group and in (B, C, E, G) from ≥ 2 biological replica with n ≥ 3 per group. Statistical significance was calculated in (B) with Brown-Forsythe/Welch ANOVA test and Dunnett’s T3 multiple comparisons test, in (C) with Kruskal-Wallis test and Dunn’s multiple comparison test, in (G) with an ordinary two-way ANOVA and Turkey’s multiple comparisons test. *p < 0.05, **p < 0.01, ***p < 0.001, ns = not significant.

**Figure S5.**
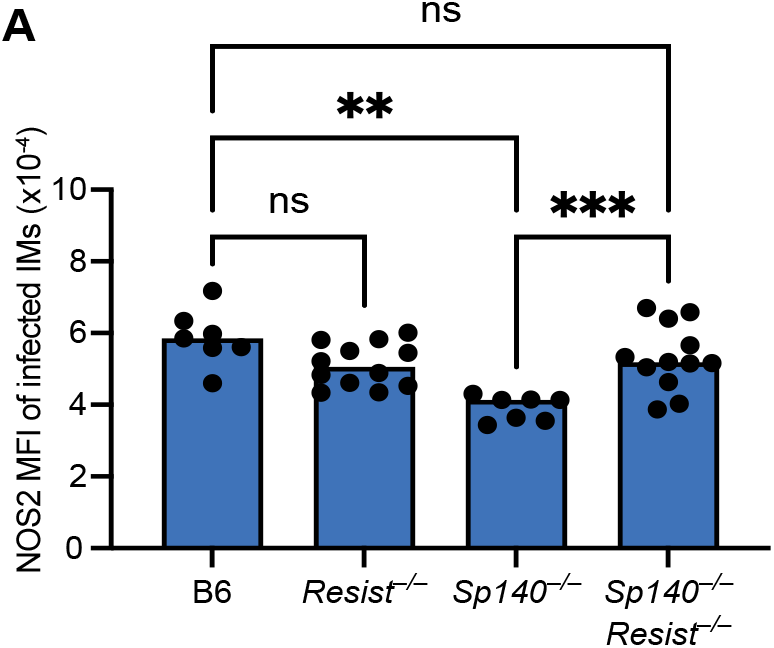
Supplementary to Figure 5. (**A**) MFI of NOS2 staining in IMs from *Mtb*-infected B6, *Resist*^*–/–*^, *Sp140*^*–/–*^, *Sp140*^*–/–*^*Resist*^*–/–*^ mice at day 26-28 pi. Statistical significance was calculated with Brown-Forsythe/Welch ANOVA test and Dunnett’s T3 multiple comparisons test. *p < 0.05, **p < 0.01, ***p < 0.001, ns = not significant.

## Notes

### Summary of Updates

Figure 1B labeling updated to correct a mislabeling error.

